# Lenacapavir allosterically remodels the HIV-1 capsid

**DOI:** 10.64898/2026.01.05.697065

**Authors:** Nayara F. B. dos Santos, Jacob A. Lewis, Mason Hansen, Miguel J. B. Pereira, Devin E. Christensen, Wesley I. Sundquist, Barbie K. Ganser-Pornillos, Owen Pornillos

## Abstract

Lenacapavir (LEN) is a highly potent, long-acting HIV-1 capsid inhibitor that holds exceptional promise for pre-exposure prophylaxis. LEN causes the mature viral capsid to rupture and lose integrity, but the underlying mechanism has been unclear. Here, we show that LEN is an allosteric modulator of HIV-1 capsid structure that breaks the capsid’s fullerene cone architecture in two steps: loss of high-curvature declinations occurs early, followed by failure of the capsid body. At the molecular level, LEN alters the non-covalent bonding interactions between capsid subunits and reduces local lattice curvature. LEN also alters the material properties of the capsid, by increasing brittleness. These results provide molecular rationales for how LEN remodels HIV-1 capsid structure and impairs the replication capacity of the virus.

## Introduction

Lenacapavir (LEN) is a highly potent, long-acting HIV-1 inhibitor (*1–3*). Initially approved for treating multidrug-resistant HIV-1, LEN has now been approved worldwide for pre-exposure prophylaxis (PrEP), holding exceptional promise with clinical trials indicating up to 100% efficacy in preventing transmission (*4, 5*). Unlike other antiretrovirals that target viral enzymes, LEN targets the capsid protein (CA), and thus interferes with multiple different stages of viral replication (*2, 3*). While LEN can bind to CA subunits in both the immature and mature virions, its highest binding affinity is for the mature assembled capsid (*1–3, 6*). Correspondingly, LEN is most potent at disrupting steps in the post-entry stage of viral replication – including nuclear import and integration (*1, 2*) – which are critically dependent on a functional, mature HIV-1 capsid.

How LEN binding to the mature HIV-1 capsid leads to potent anti-viral activity is not fully understood. The HIV-1 capsid is a fullerene cone protein shell that organizes and protects the viral genome (*7*), and is the principal viral interface for engaging intracellular host factors (*8*). LEN shares binding sites on the capsid surface with FG-motif (phenylalanine-glycine) containing host proteins (*9, 10*), but competition with host factor binding does not fully explain the drug’s potency or mechanism of action (*3, 11, 12*). Rather, LEN appears to work by altering capsid architecture. Although initially reported to stabilize intact capsids (*3*), subsequent studies have shown that LEN treatment causes capsids to lose integrity in an apparently dynamic process: initially, the capsid ruptures and loses some fraction of its CA subunits; ultimately, the remaining CA subunits persist in the form of a hyperstable, assembled capsid fragment (*11, 12*). The high-resolution structural details of these changes – and their consequences for global capsid integrity – have not been fully clear.

In this study, we used electron microscopy imaging to analyze the structures of purified HIV-1 cores treated with LEN. We found that initial capsid rupture occurs in regions containing CA pentamers, and subsequent damage occurs in the main body of the cone. The resulting capsid remnants are missing their declinations and contain long fissures that separate flattened lattice segments composed of LEN-bound CA hexamers. Focused reconstructions indicate that LEN binding alters inter-subunit contacts that are distal from the binding site, implying an allosteric mode of action. Our results reveal the high-resolution details of how LEN reshapes the HIV-1 capsid, and provide new insights on the molecular basis of its antiviral activity.

### LEN fractures the HIV-1 capsid

Previous high-resolution structural studies of LEN-capsid interactions have relied only on soluble, disulfide-stabilized HIV-1 CA hexamers or in vitro CA assemblies (*2, 3*). In contrast, we performed our analysis on actual viral capsids that were purified from virions (**Fig. S1A-D**). Initially, we examined how LEN affects gross capsid morphology, at pharmacologically relevant drug concentrations (protocol schematic in **Fig. S1E**). Purified cores (1 nM of CA) were incubated with 5 nM LEN, which is the EC95 of the drug and approximates the drug plasma concentration at 24 weeks after subcutaneous delivery (*2*). Cores were sampled at 0, 1, 4 and 8 h of incubation, and then analyzed by using negative stain electron microscopy.

After 1 h of incubation with LEN, essentially all capsids had already fractured, as evidenced by clear gaps in projection images of the capsid walls (**Fig. 1A,B**); interestingly, these early gaps were located at or near the pentamer-containing declinations (indicated by white arrowheads in **Fig. 1A**). At longer incubation times (4 and 8 h), the capsids had even more fractures and wider gaps (**Fig. 1A,B**). The capsid surfaces also increasingly flattened (**Fig. 1A,C**). In contrast, >90% of cores retained apparently intact capsids at all time points when incubated with DMSO control (**Fig. 1A-C**). These results confirm previous studies (*11, 12*) and indicate that, at pharmacologically relevant concentrations, LEN binding causes physical damage to the HIV-1 capsid, at least in vitro.

**Figure 1.**
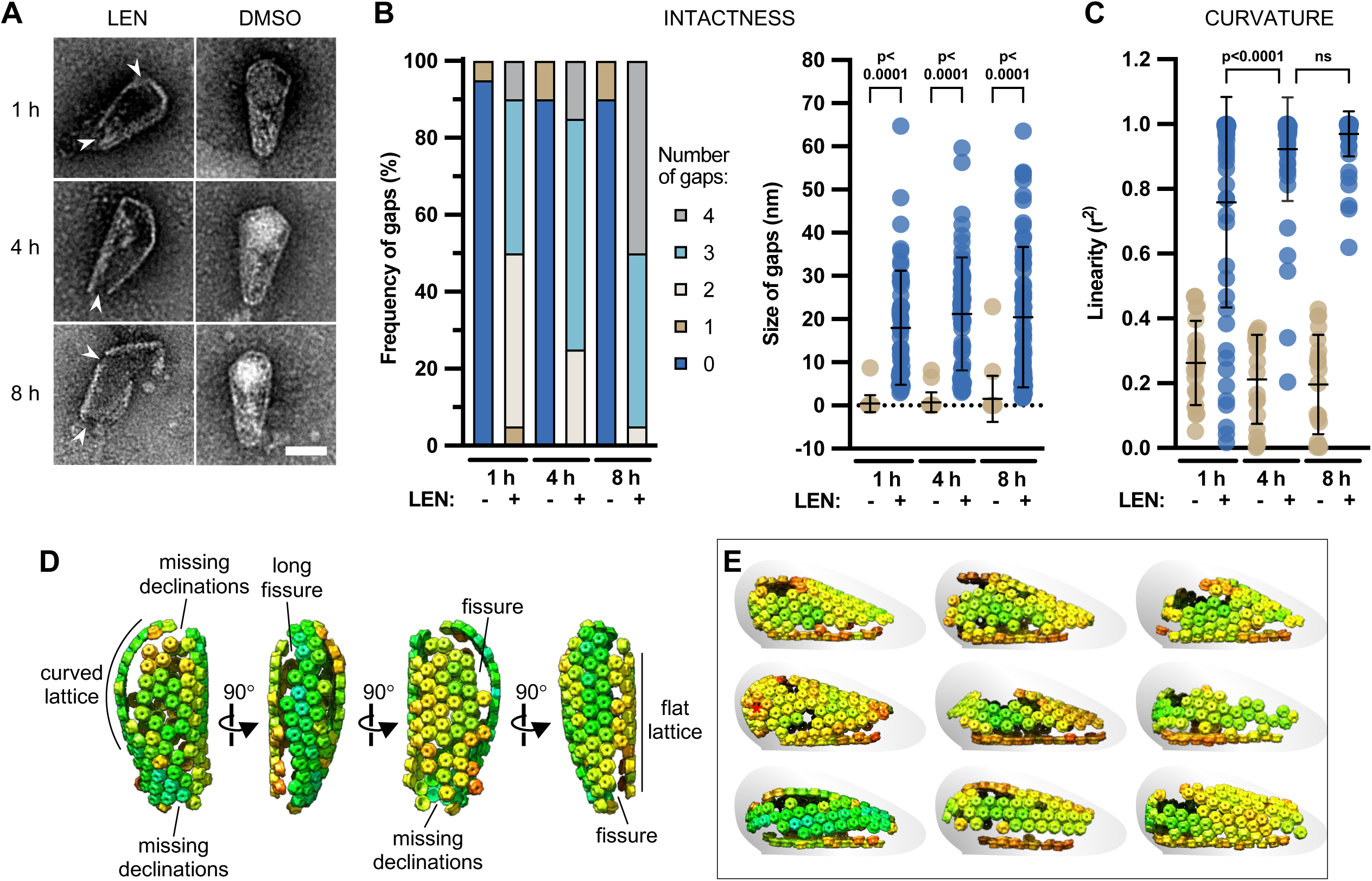
LEN fractures HIV-1 capsids. (**A**) Negative stain images of representative viral cores after incubation with 5 nM LEN (left column) or DMSO control (right column) for the indicated number of hours. Arrowheads point to missing declinations. (**B**) Quantification of fracturing (n = 20 randomly selected cores). Left panel, number of gaps observed per capsid. Right panel, sizes of the gaps. (**C**) Quantification of curvature for the same cores. (**D**) Orthogonal views of an illustrative capsid remnant, after incubation with excess LEN to drive fracturing to completion. Lattice map is shown as hexamers colored according to cross-correlation values from sub-tomogram averaging, with red showing low correlations. (**E**) Gallery of cores that exemplify shared fracturing characteristics. Capsid remnants are oriented with the flat wall at the bottom. Red asterisk (left column, middle row) indicates the inferred position of an intact pentamer.

Next, we examined cores incubated with excess LEN, to gain additional insights on the nature of the capsid remnants at the end point of LEN incubation. In this experiment, core concentrations were sufficiently high (5-10 μM CA) to allow imaging by cryoEM. Importantly, these capsid remnants recapitulated the end point characteristics of cores incubated with 5 nM LEN (**Fig. S2A-C**), indicating that the only difference between the high and low concentration regimes was the kinetics of fracturing.

We used electron cryotomography (ECT) and sub-tomogram averaging lattice mapping to visualize the three-dimensional organization of the fractured capsids (**Fig. 1D-E**, **Fig. S3A-C**). Because this approach tends to underestimate completeness, we initially restricted our analysis to cores whose lattice maps clearly outlined the cone shape and that retained a vast majority of their subunits (**Fig. 1E**). Interestingly, we found that what can appear to be multiple, separate fragments in projection images actually comprise a single capsid remnant (**Fig. 1D**). The most complete lattice maps confirmed that declinations were largely missing, although individual pentamers could still remain (**Fig. 1E**, asterisk). Most notably, the capsid bodies were split by one or more fissures that could run along the entire length of the cone. The fissures delineated opposing walls of distinct arcs – flat or curved – giving the capsid remnant the appearance of a damaged shoe. In a few cases, both the top and the bottom of the shoe were flattened, with curved sides.

Having identified the key characteristics of the LEN-damaged capsids, we were then able to analyze the full dataset (71 cores total). Although 8/71 had too limited coverage to analyze, 63/71 (89%) could be classified by visual inspection as being incomplete lattice maps of similar-looking capsid remnants (**Fig. S3C**). These findings indicate that, despite the extremely high degree of pleomorphism in HIV-1 capsids, LEN breaks them all through the same mechanism.

We also confirmed that LEN induces the same morphological changes in synthetic capsids obtained by assembling purified recombinant CA protein (**Fig. S2D-F**). These results indicate that the structural effects of LEN on the capsid are independent of the contents of the core. The synthetic capsids were more resistant to LEN effects (more below), which we believe is due to inherently lower strain in the empty capsids (*13*).

Together, our results support findings that, rather than having a globally stabilizing effect on the HIV-1 capsid, LEN disrupts the fullerene cone architecture (*11, 12*). Importantly, two types of fractures form at pharmacologically relevant LEN concentrations: 1) loss of declinations at earlier time points, and 2) fissuring of the capsid walls at later time points.

### LEN alters the CA hexamer structure

To determine the molecular mechanism through which LEN alters capsid morphology, we used single-particle averaging cryoEM approaches (*14, 15*) to examine the structures of the CA capsomers from LEN-treated and untreated cores. Focused reconstruction indicated that the LEN-saturated cores primarily contained hexamers, whose average structure we solved to a nominal resolution of 2.8 Å (**Fig. 2A, Fig. S4A-B**). The drug density in the asymmetric unit (**Fig. 2A**, inset) has the same definition as the surrounding sidechains, confirming full occupancy of all six possible binding sites in the hexamer. We also solved the average CA hexamer structure from unbound cores, to a nominal resolution of 2.9 Å (**Fig. S5A,C**). Coordinate models were refined into each map, to allow comparison of the two structures in molecular detail (**Table 1**).

**Figure 2.**
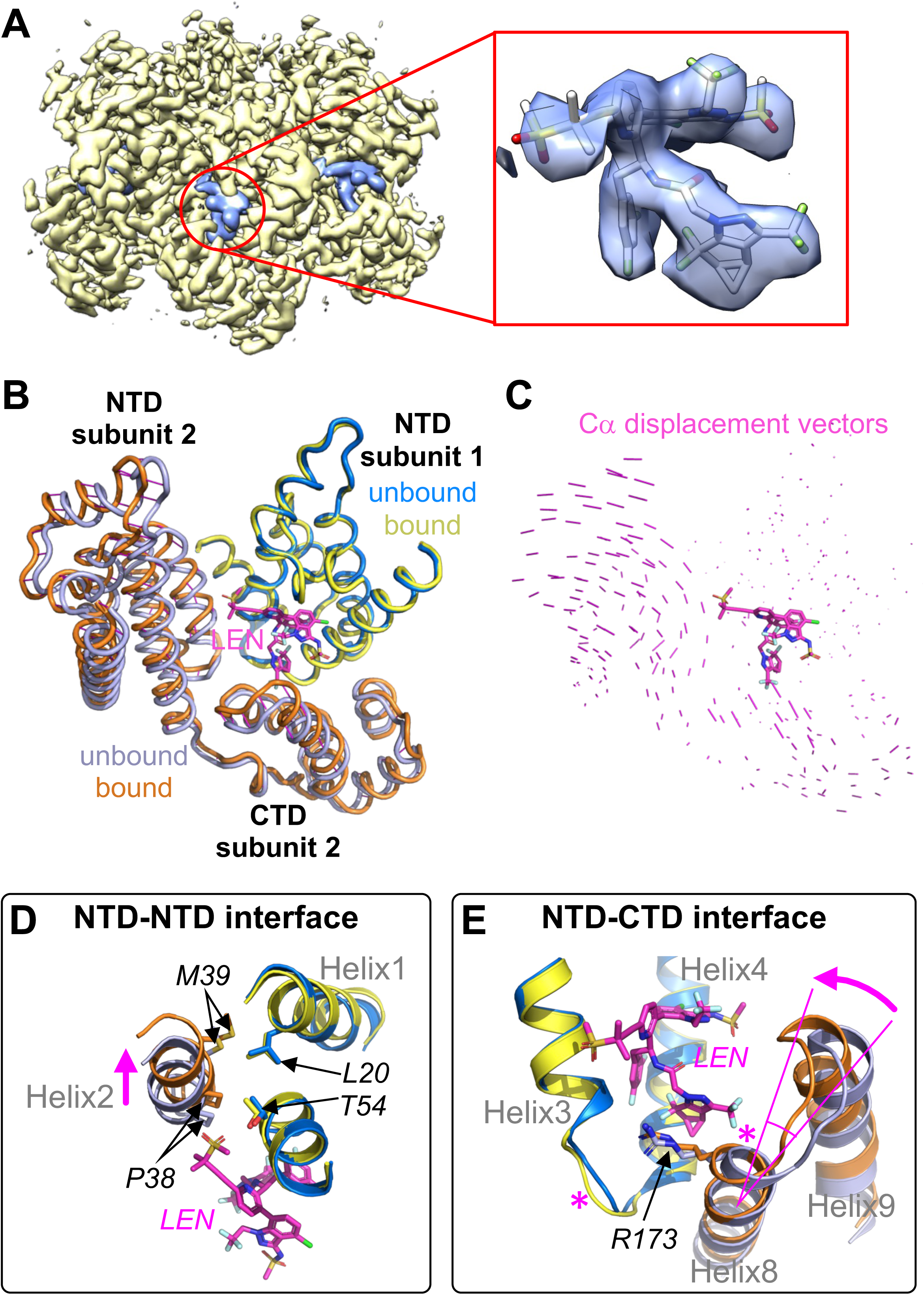
Comparison of LEN-bound and unbound CA hexamers from purified HIV-1 cores. (**A**) CryoEM density map of LEN-bound hexamer at 2.8 Å nominal resolution, with protein densities colored in yellow and LEN densities in light blue. Inset (red box) shows a close-up of the drug density in the asymmetric unit, together with the final refined coordinate model. (**B**) Comparison of the repeating interactions that form the hexamer from LEN-bound and unbound structures, which are superimposed on the NTD of one subunit. LEN is shown in sticks representation (magenta). (**C**) Displacement vectors, indicating the extent to which each subunit differs in position. LEN is shown in sticks for reference. (**D,E**) Details of the NTD-NTD interface (D) and NTD-CTD interface (E). LEN, helices, and landmark residues are labeled. Magenta arrows indicate shifts in position induced by LEN binding. Asterisks indicate conformational changes.

**Table 1.** Image data collection and map reconstructions.

| Sample | Saturated Cores |  | Unbound Cores |  |  | Unsaturated Cores | Unsaturated In Vitro Capsids |  |  |  |
| --- | --- | --- | --- | --- | --- | --- | --- | --- | --- | --- |
| Magnification | ×81,000 |  | ×81,000 |  |  | ×81,000 | ×81,000 |  |  |  |
| Voltage (kV) | 300 |  | 300 |  |  | 300 | 300 |  |  |  |
| Dose (e-/Å<sup>2</sup>) | 50 |  | 50 |  |  | 50 | 50 |  |  |  |
| Defocus range (μm) | 0.5 to 2.5 |  | 0.5 to 2.5 |  |  | 0.5 to 2.5 | 0.5 to 2.5 |  |  |  |
| Pixel size (Å) | 1.06 |  | 1.08 |  |  | 1.08 | 1.06 |  |  |  |
| Focus Region | Hexamer | Hexamer-hexamer interface | Hexamer | Hexamer | Hexamer-hexamer interface | Pentamer | Pseudo 3fold | Pentamer | Hexamer next to pentamer | Hexamer-hexamer interface |
| EMDB | 72657 | 73410 | 71816 | 71817 | 73468 | 73397 | 73398 | 73399 | 73400 | 73401 |
| Symmetry imposed | C6 | C2 | C1 | C6 | C2 | C5 | C1 | C5 | C1 | C2 |
| Particles | 1,064,761 | 1,064,761 | 901,622 | 901,622 | 384,735 | 119,349 | 877,757 | 518,565 | 1,669,740 | 740,861 |
| Resolution at FSC 0.143 (Å) | 2.8 | 3.2 | 3.3 | 2.9 | 3.4 | 4.5 | 3.5 | 3.5 | 3.4 | 3.8 |
| Map Resolution Range (Å) | 2.3 to 34 | 2.7 to 40 | 2.4 to 35 | 2.4 to 24 | 3.0 to 48 | 2.5 to 64 | 2.3 to 29 | 2.3 to 7.9 | 2.3 to 8.2 | 3.2 to 41 |
| Map Sharpening B-factor (Å<sup>2</sup>) | 117.1 | 122.6 | 100.1 | 108.4 | 120.0 | 153.6 | 117.2 | 132.0 | 143.7 | 165.4 |

Superposition of the unbound and LEN-bound models as full hexamers revealed that they are very similar, with average root-mean-squared-deviation over equivalent Cα atoms of 1.5 Å (**Fig. S6A**). Importantly, however, the displacement vectors, although small, had non-random trajectories. This result indicated that LEN induces small, but coordinated changes in the CA hexamer architecture, which become more evident when superpositions are done at the level of individual subunits (**Fig. 2B,C**, superimposed on NTD of subunit 1; see also **Fig. S6B**). Through systematic analysis of such superpositions, we found that LEN alters packing of CA subunits at each of the two intermolecular contacts that stabilize the hexamer, which are called the NTD-NTD and NTD-CTD interfaces (*16*). At the NTD-NTD interface, packing of helix 2 against helices 1 and 3 is altered by a slight rotation and translation of helix 2 (**Fig. 2D**, magenta arrow). At the NTD-CTD interface, the orientation of the CTD relative to the NTD is altered by ∼8° (**Fig. 2E**, magenta arrow), along with small changes at the N-terminal end of helix 8 and the loop connecting helices 3 and 4 (**Fig. 2E**, asterisks). LEN also altered the configuration of the N-terminal β-hairpin (**Fig. S6C**). Crucially, these changes are distal (15-30 Å away) from the LEN binding site, and thus identify LEN as an allosteric modulator of the HIV-1 CA hexamer.

### LEN reduces local lattice curvature

LEN binds at the NTD-CTD interface, which is a crucial determinant of lattice curvature (*16–19*). To learn how LEN affects lattice curvature, we performed focused reconstructions centered on two adjacent hexamers (**Fig. S4**, **Fig. S5**). In these maps, two drug binding sites flank the CTD-CTD dimer interface that connects the two hexamers. In comparing the LEN-bound and unbound maps, we found that LEN induces rotation of the NTDs toward the CTD, and thus the flanking NTDs tilt towards each other (**Fig. 3A,B**). In CA tubes, LEN binding is reported to stabilize the CTD-CTD dimer that connects the hexamers (*3*), but the dimer structure is essentially unchanged in comparisons of our bound and unbound maps (**Fig. 3C,D**). Unlike in CA tubes, the CTD dimer shows little variation in actual capsids, since flexion of the NTD-CTD interface is the principal determinant of lattice curvature in the capsid body (*15, 19*).

**Figure 3.**
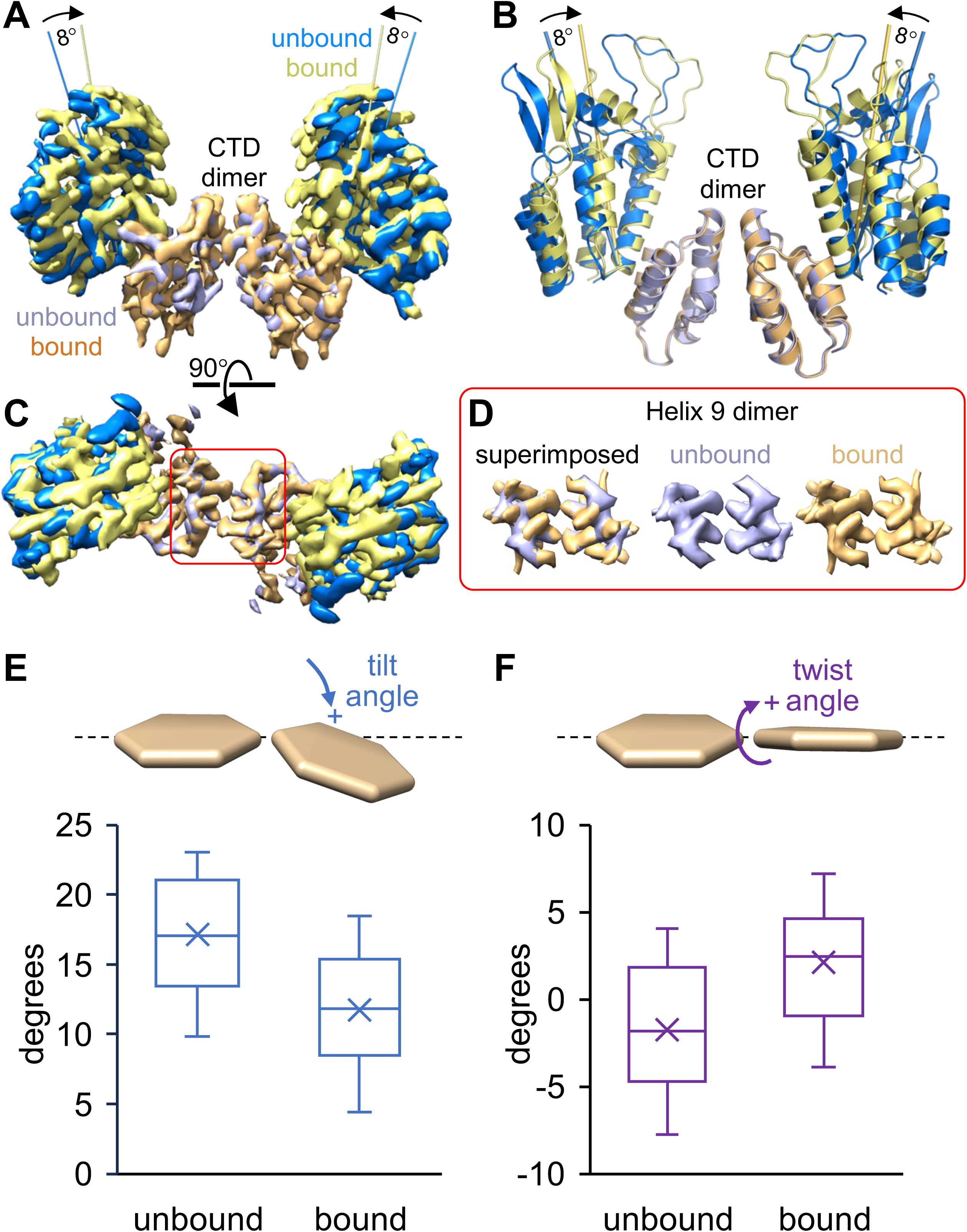
Effect of LEN on local lattice curvature. (**A**) Side view of superimposed cryoEM maps of NTD-CTD interfaces from two hexamers connected by the CTD-CTD dimer. The unbound (3.5 Å) and bound (3.5 Å) maps were superimposed on the CTD only, revealing differing orientations of the NTDs. (**B**) Rigid body docked models into the superimposed maps. (**C**) Top view of the maps. (**D**) Cropped views of the helix 9 dimer. (**E,F**) Definitions and box-and-whisker plots of hexamer-hexamer tilt (E) and twist (F). Horizontal lines indicate the quartiles, × indicates the mean of measured angles from 15 maps.

Our consensus maps describe the overall average curvature in the bound and unbound capsids. To quantify how LEN affects curvature more precisely, we defined particle subsets that each spans a narrower range of curvatures (*15*). We used these subsets to reconstruct 15 maps and then quantified the relative orientations of the two hexamers in each map, in terms of tilting and twisting angles (**Fig. S7**). We found that the average hexamer-hexamer tilt is reduced by 5.4° in the LEN-bound structures (**Fig. 3E**), consistent with relative flattening of the capsid surface. There is also a shift to more positive twist in LEN-bound capsids (**Fig. 3F**). Importantly, these results also indicate that LEN-induced allosteric changes in the CA hexamer can propagate throughout the capsid lattice.

Previous studies have suggested that curvature in the HIV-1 capsid is primarily due to an asymmetric (or warped) CA hexamer (*20*). We explored this possibility by using the warped hexamer as initial template and not imposing symmetry in reconstructing the unbound hexamer, but this approach still yielded a closely 6-fold symmetric map (**Fig. S5A,B**). Thus, we conclude that LEN-induced curvature effects need not invoke a mechanism of dramatic symmetry change within the hexamer ring itself.

### LEN occupancy correlates with hexamer-hexamer tilt

Next, we examined the molecular basis of the early fracture events that lead to loss of the declinations; these sites contain the CA pentamers, as well as hexamers with the highest local curvatures. The early effect is intriguing, because LEN is predicted to bind only to CA hexamers and not pentamers (*14, 15, 21*). LEN-saturated cores contained virtually no pentamers, and so we incubated cores with sub-saturating levels of drug (4:1 ratio of CA to LEN). In this case, we were able to perform focused reconstruction on the CA pentamer, but unfortunately could not achieve high resolution (**Fig. S8**). To reach higher resolution, we prepared an equivalent sample with in vitro assembled capsids and obtained reconstructions centered on the pentamer (3.5 Å) and on the hexamer adjacent to the pentamer (3.4 Å) (**Fig. S9**). Both maps confirmed that the pentamer does not bind to LEN. The fact that we were able to solve the pentamer to higher resolution from empty capsids but not cores indicates that cores are more susceptible to LEN rupturing, presumably because the core contents induce capsid strain (*13*).

The six binding sites on the hexamer next to the pentamer had differing LEN occupancies (numbered in clockwise direction, **Fig. 4A**): the two sites closest to the pentamer (sites 1 and 2) had high and low occupancies, respectively; the next adjacent two sites (3 and 6) were also high, whereas the most distal sites (4 and 5) were intermediate (**Fig. 4A-B**). We further confirmed this high-low-high-intermediate-intermediate-high pattern by reconstructing a map centered on the pseudo 3-fold of two hexamers and one pentamer, and examining the two distinct hexamers in this map (**Fig. S10**). Thus, the LEN binding pockets on the capsid surface are not equivalently occupied by LEN at sub-saturating drug levels, at least in hexamers next to pentamers.

**Figure 4.**
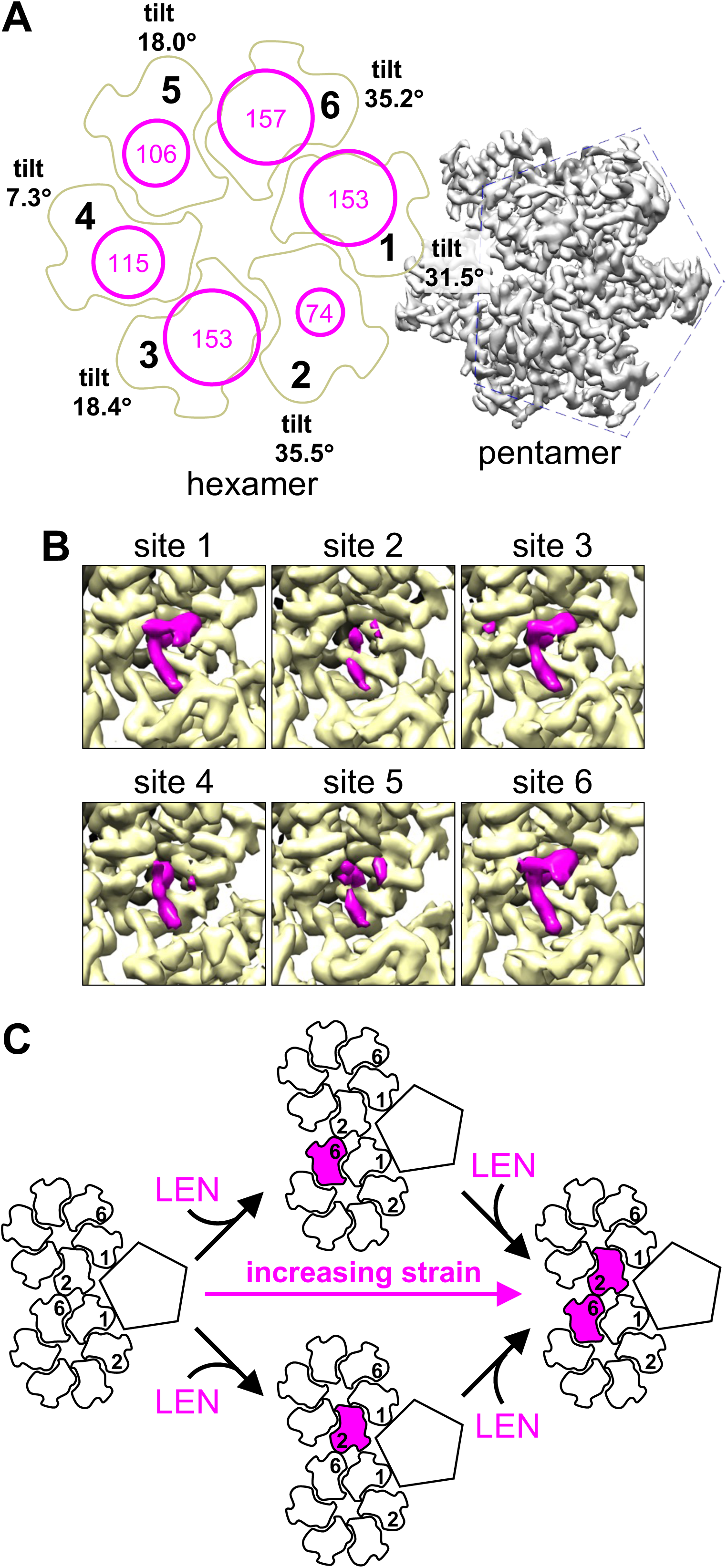
Effect of LEN on the pentamer-containing declinations. (**A**) Schematic of the hexamer next to a pentamer. Subunits are numbered 1 through 6 to distinguish the LEN binding sites; site 1 connects to the pentamer. Magenta circles indicate relative LEN occupancy in our consensus map centered on the hexamer; radius is proportional to occupancy. Occupancy values (magenta) are expressed as volume of the LEN density (Å^3^) when measured at map contour level of 0.34. Local tilt angles are also indicated. (**B**) Close-up views of LEN densities (magenta) at the six sites, with map contoured at 0.34. (**C**) Schematic model depicting how LEN binding to sites 2 and 6, which flank the highest tilt hexamer-hexamer interface in the entire fullerene cone, induces local strain.

The hexamer next to the pentamer has the highest local tilts in the entire fullerene cone, and also can have the greatest variation in this angle across its six subunits (**Fig. 4A**). Accordingly, we checked for correlations between LEN occupancy and tilt. Since LEN reduces local curvature, we were intrigued to find that sites with highest tilt (35°) had either the highest (site 6) or lowest (site 2) mean occupancies. Importantly, these two sites flank the highest tilt hexamer-hexamer interface in the capsid (**Fig. 4C**). Site 1 also has high tilt (31°) and high occupancy, but this site faces the pentamer which cannot bind LEN (**Fig. 4A-C**). Our interpretation is that high tilt interfaces tolerate LEN at one of the two flanking binding sites, but simultaneous occupancy of both sites is likely to be destabilizing (**Fig. 4C**).

Given these observations, we checked whether tilt and occupancy also correlate in hexamers located in the body of the cone. From a consensus map focused on two hexamers (3.8 Å) (**Fig. S11A-B**), we sorted particles according to tilt, and then reconstructed high tilt and low tilt maps (both 4.1 Å; **Fig. S11C-D**) to compare mean occupancies. In this case, we found that the low tilt site (**Fig. S11E**) had substantially higher mean occupancies than high tilt site (**Fig. S11F**). Unfortunately, we were unsuccessful in doing the complementary analysis – sorting particles by occupancy in order to compare tilts – because the drug mass is too small compared to the total mass in our maps.

### LEN anti-viral potency changes with time of addition

The above results suggested to us that LEN’s potency as an antiviral would depend on how long the virus is exposed to the drug. We tested this by using a single-cycle infectivity assay. We transduced THP-1 cells with HIV-GFP reporter virus, and then tested the effects of LEN when added at different times post-transduction (**Fig. 5A**). We first performed a reference experiment at high LEN concentration: 250 nM LEN was completely inhibitory when added as late as 4 h post-transduction, but was no longer effective when added 8 h post-transduction or later in this assay system (**Fig. 5B**). Since 250 nM LEN completely fractures capsids within minutes (*11*), the loss of activity at the 8 h time point indicates that the capsid is no longer accessible to the drug or has already completed its essential functions. In any case, this defines the time window (i.e., 0-4 h post-transduction) in which to interpret results at low LEN concentrations. We then performed the same experiment at 5 nM LEN (**Fig. 5C**). In this case, 15% infectivity was observed when LEN was added at transduction, with progressively higher infectivity when added 1 h (40%) or 4 h (86%) post-transduction. We also measured the apparent EC50s, and indeed, these values progressively increased with later time points of drug treatment (**Fig. 5D**). These results support the conclusion that low LEN concentrations require longer exposure to the capsid to achieve their full effect (*11*).

**Figure 5.**
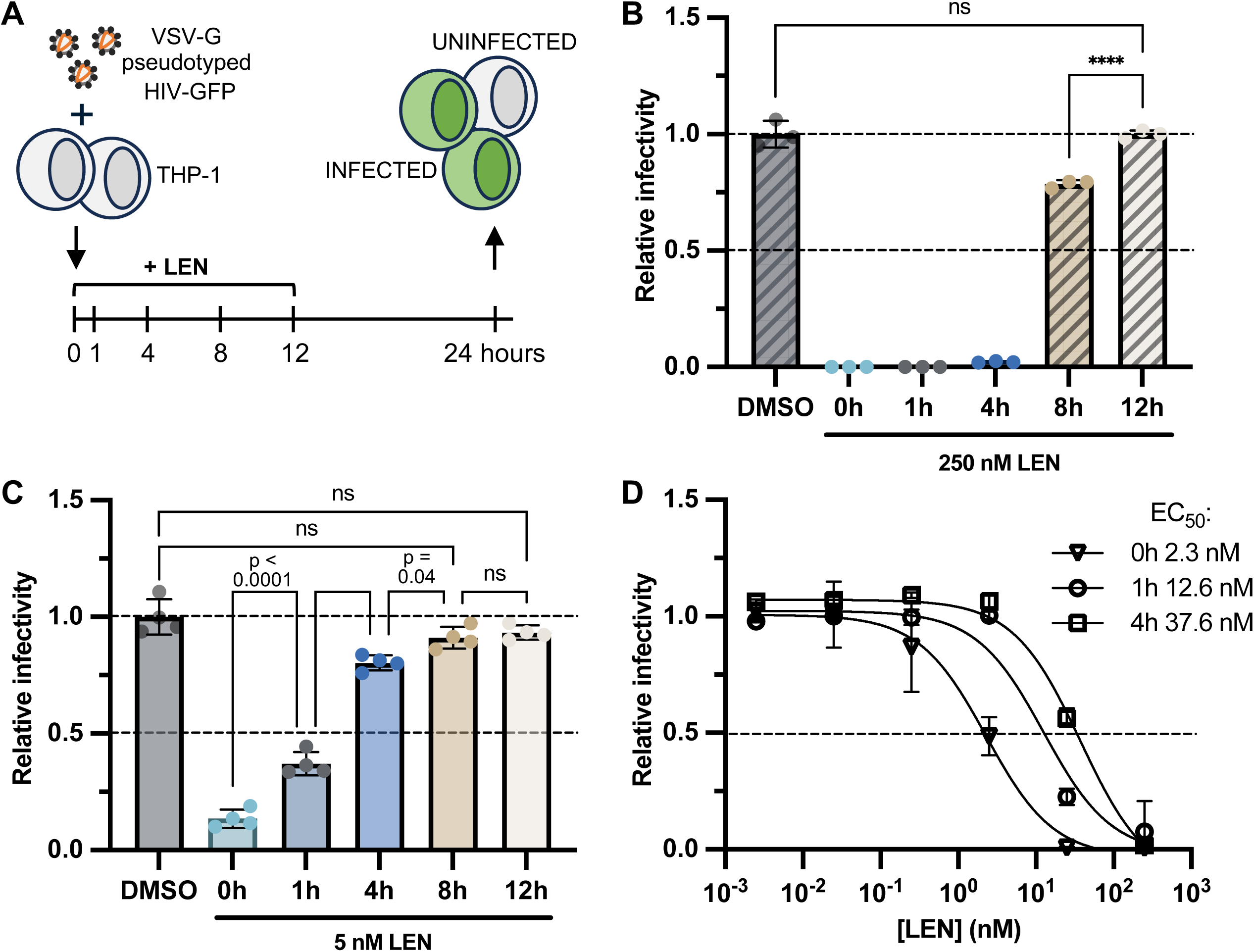
LEN potency depends on time of addition in a single cycle infectivity assay. **(A)** Experimental schematic. (**B**) Effect of 250 nM LEN on infectivity of VSV-pseudotyped HIV-GFP reporter virus at 25 nM CA. (**C**) Effect of 5 nM LEN. **(D)** Apparent EC50s at indicated times of addition, as determined by curve fitting. Graphs depict mean ± s.d. from three or four technical replicates, representative of two or three independent experiments. Statistics were calculated using two-way ANOVA with Dunnett’s post hoc test.

## Discussion

Here, we show that LEN is an allosteric modulator of HIV-1 capsid structure that alters the non-covalent bonding interactions between the CA subunits at near-atomic resolution scale, and fractures the fullerene cone at nanometer scale. We can rationalize these observations as a form of stress-strain response (**Fig. 6**). We propose that LEN binding induces stress on the capsid lattice, and in response, the assembled CA subunits adjust to reduce strain. As more and more LEN molecules bind, strain accumulates to the point of material failure, which is when the capsid fractures. At high (μM) LEN concentrations, binding occupancy and stress rise too fast to be accommodated, leading to almost immediate failure (**Fig. 6A**). On the other hand, we observed two types of failure at low (5 nM) LEN: loss of declinations within 1 h (the earliest time point in our experiments), and fissuring of the cone body within 4 h. This interesting finding indicates to us that the fullerene cone does not have uniform material properties. Rather, the declinations and capsid body respond differently to stress and have different failure thresholds (**Fig. 6B**). In this model, the declinations have lower threshold and thus fail first, which likely also relieves overall strain on the remaining capsid. The body of the cone can tolerate higher strain, and thus requires higher LEN occupancy to reach its failure threshold. In agreement with our model, in silico simulations also indicate that the fullerene cone has a material heterogeneity, with the ends of the cone (where the declinations are located) having higher stiffness than the cone body (*22*). All of these effects are a consequence of the fullerene cone architecture, in which every single capsid subunit occupies a unique environment.

**Figure 6.**
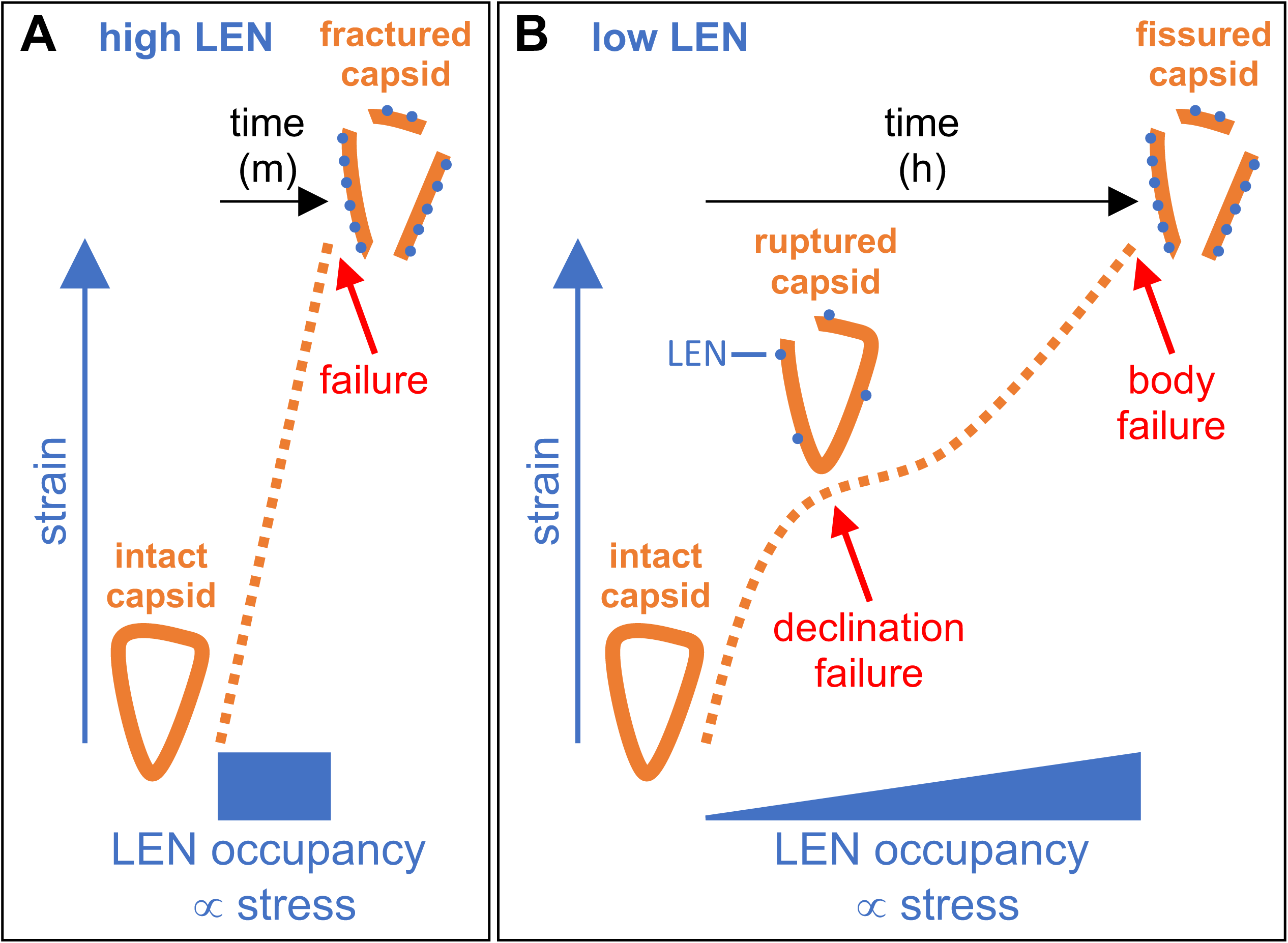
Stress-strain model of LEN-induced capsid fracturing. The general scheme is that strain (vertical axis) increases as a function of applied stress (horizontal axis). In this case, LEN applies stress on the fullerene cone in proportion with binding occupancy. (**A**) At high LEN concentrations, occupancy and stress rise too fast to be accommodated, leading to almost immediate material failure (red arrow). (**B**) At low LEN concentrations, occupancy and stress rise more slowly, and thus the differing failure thresholds of the declinations and main cone body can manifest (red arrows). Since we observed a few intact pentamer positions in our lattice maps, it seems likely that different declinations will also have different fracture thresholds, in which case the stress-strain curve will not be as smooth as depicted here.

Interestingly, single-molecule experiments have shown that LEN occupancy at the time of capsid rupture increases with LEN concentration (*11*). At low LEN, rupture occurs at 20–30% mean occupancy, which we now interpret to correspond to declination failure. At high LEN, occupancy rises to ∼70% at the time of capsid breakage (*11*), which we interpret to correspond to failure of the capsid body. The lower fracture threshold of declinations is likely to be relevant to uncoating, the virus replication step in which the capsid opens to release its genome.

Our finding that LEN allosterically modulates capsid structure is consistent with the fact that LEN binds at the juncture of major CA-CA interactions, a location that is critical in determining curvature of the hexagonal capsid lattice (*16–19*), hexamer/pentamer switching (*14, 15, 23*), and host factor binding (*9, 10, 24–28*). Our analysis of LEN binding to high tilt hexamers suggests to us a molecular rationale for how binding of more and more LEN molecules eventually leads to fracturing of the capsid. Key to this model is the relative occupancy of the two sites that flank high tilt hexamer-hexamer interfaces (**Fig. 4C**). We propose that LEN binding to high tilt sites induces more local strain than binding to low tilt sites, and the fracture threshold is therefore determined by the number and proximity of drug-bound high tilt sites. Declinations cluster high tilt sites at the ends of the cone, which are thus more prone to fracturing than the capsid body (*22, 29*).

Another crucial finding from our studies is that LEN fundamentally alters the non-covalent bonding interactions that hold together the capsid. Whereas these alterations can be rationalized as simply the basis of LEN’s well-established stabilization of the hexagonal CA lattice (*1, 3, 11, 12*), we propose the broader implication that they also change the properties of the capsid as a material. A possible analogy is alloying in metallurgy, the ancient practice of mixing another element with a base metal. In an alloy, the introduced atoms typically alter metallic bonds and can create internal stresses that change the properties of the material and, importantly, how it responds to environmental stressors (*30*). In support of this idea, our lattice maps of the LEN-treated capsids show fractures with well-defined edges, which is indicative of the material property of brittleness (*31*). Capsids treated with GS-CA1 (a LEN analog) during reverse transcription also show brittle fractures (*32*). In contrast, lattice maps of ruptured capsids during reverse transcription but in the absence of drug show more plastic deformation (i.e., they are less brittle) (*13, 32*).

LEN dose response curves in single-cycle assays are biphasic (*11*), as are those of PF74 (*9, 33*), a related but much less potent compound. A low concentration phase (pM for LEN) inhibits infection up to 10-fold, followed by a high concentration phase (nM for LEN) that tracks with inhibition of reverse transcription. Our structural observations are most easily correlated with the high concentration phase, in which LEN inhibits reverse transcription by damaging the capsid (*11, 32*). Consistent with this interpretation, preincubation increases drug potency in the second phase (*11*); conversely, late addition decreases potency as we find here.

The molecular basis of drug activity at the first phase, where reverse transcripts can accumulate to normal levels, remains to be explained. In vitro experiments show that LEN can still rupture capsids in this concentration regime, although occupancy builds up with much slower kinetics (*11*). We propose that, even at occupancies where LEN by itself is insufficient to break the capsid, the rupture threshold can still be reached when augmented by other stressors. This additive effect has already been observed in two contexts. Firstly, simulations show that the packaged contents of the core induce capsid strain (*13*), and we find here that cores are more prone to LEN-induced fracturing than empty capsids. Secondly, reverse transcription stresses the capsid from within (*29, 34*), and GS-CA1 dramatically exacerbates capsid rupture during this process (*32*).

Capsid brittleness is implicated in HIV-1 nuclear import (*22, 35, 36*), a replication step that LEN inhibits in the low concentration regime (*1, 2*). HIV-1 CA mutants that are impaired in this step form brittle (inelastic) capsids, and compensatory mutations that restore import also restore elasticity (*22, 37*). According to our model, LEN-induced brittleness should make the capsid prone to rupture as it is subjected to the stresses of passaging through the nuclear pore, and indeed, simulations support this prediction (*38*). Thus, the interconnected nature of the capsid lattice, its unique fullerene cone architecture, its mechanical properties, and the stresses associated with replication and nuclear entry, combine to make the capsid susceptible to allosteric inhibition by LEN.

## Materials and methods

### Preparation of HIV-1 cores

One day before transfection, 5.5 × 10^6^ of HEK293FT cells (American Type Culture Collection) were seeded in 15-cm dishes in DMEM supplemented with 1× MEM, 100 mM HEPES, Pen-Strep, and 10% FBS, then incubated overnight at 37 °C in 5% CO_2_. Cells were transfected with 24 μg of pSG3^ΔEnv^ plasmid using the calcium-phosphate method (Takara). After 24 h, the media was replaced with exosome-free media (*32*) (prepared by overnight ultracentrifugation in a Beckman SW-32 rotor at 28,000 rpm at 4 °C, followed by dilution of the supernatant in DMEM down to 10% (*v*/*v*) FBS). Cells were then incubated for an additional 72 h. The supernatant was collected and treated with 175 μg/mL subtilisin (Sigma) and 7 U/mL of benzonase (Sigma) at 37 °C for 1 h. Subtilisin was inactivated with 4.35 μg/mL PMSF for 15 min at room temperature. Virions were pelleted by ultracentrifugation through a 20% (*w*/*v*) sucrose cushion at 28,000 rpm in an SW-32 rotor for 1 h at 4 °C. Pellets were resuspended in virus buffer (50 mM Tris, pH 8, 85 mM NaCl). To lyse the viral membrane, virions were incubated with 190 μM IP_6_ (phytic acid, Sigma) and 0.35 mg/mL of melittin (Sigma) for 10 min at 37 °C. To release cores from residual membranes, the sample was treated with 0.45 mg/mL digitonin for 2 min at room temperature, followed by tabletop centrifugation at 17,000*g* for 20 s. The supernatant was loaded on a Superose 6 Increase 10/300 GL column (Cytiva) and eluted with buffer (20 mM Tris, pH 8, 100 mM NaCl, 80 μM IP_6_). Fractions were combined, supplemented with 10 mM ATP (Promega) and concentrated using a 30-kDa cutoff Amicon Ultra-4 concentrator (cellulose acetate filter). Yields were quantified (in terms of CA concentration) by densitometry of Coomassie-stained SDS-PAGE gels with purified SG3 CA protein as standard.

### Analysis of capsid morphology at 5 nM LEN

Purified cores were diluted to 1 nM CA and incubated with 5 nM LEN (diluted from DMSO stock) in 500 mL of 100 mM Tris, pH 8, 100 mM NaCl, 10 mM MgATP. Samples were incubated at room temperature with gentle rocking. Control samples were prepared in DMSO without LEN. After indicated incubation times (**Fig. 1A**), 4 mL of pre-equilibrated Q Sepharose resin (Cytiva) was added. After 30 min, the resin was recovered via gravity flow columns. Cores were eluted after incubation for 5 min with 2 mL of elution buffer (50 mM Tris, pH 8, 1 M NaCl, 10 mM ATP), then reconcentrated with a 30-kDa cutoff Amicon Ultra-4 concentrator. Final samples (5 μL) were applied onto Formvar/Carbon-coated, 300-mesh copper grids (Electron Microscopy Sciences) for 2 min, stained for 30 s by floating on a drop of 4% (*w*/*v*) uranyl acetate, blotted, re-stained for 1 min, and blotted dry. Images were taken with a JEM-1400 transmission electron microscope (JEOL) operating at 120 kV.

Projection images were analyzed and quantified for capsid fracturing and curvature using Fiji software (*39*). For measurements, around 20 cores were randomly selected for each condition, using these criteria: isolated, long axis parallel to the grid surface, not obscured by debris or staining artifacts. Gaps were identified as interruptions in the capsid lattice and measured using the straight-line tool. For curvature analysis, each contiguous capsid wall was traced using the segmented-line tool. Cartesian coordinates (x, y) were extracted and fit using linear regression, and the r^2^ value was taken as a measure of linearity (r^2^ = 1 is a straight line). For apparently fragmented capsids, each fragment was measured separately.

### Analysis of capsid morphology at excess LEN

Purified cores were diluted to 5 μM CA, then incubated with drug (at CA:LEN molar ratios of 1:0, 4:1, and 1:10) for 1 h prior to vitrification in C-Flat 2/1 300-mesh copper grids (Electron Microscopy Sciences) or Quantifoil R2/1 300-mesh copper grids (Electron Microscopy Sciences). Control samples were treated with equivalent volumes of DMSO.

Empty capsids were assembled by incubating 500 μM of recombinant CA protein with 100 mM MES, pH 6, 4 mM IP_6_ at 37 °C for at least 1 h. The capsids were separated from unassembled CA by using a Superdex 200 10/300 GL column (Cytiva), in running buffer (20 mM Tris, pH 8, 100 mM NaCl, 10 mM β-mercaptoethanol, 40 μM IP_6_). Fractions were combined and buffer exchanged four times, using a 30 kDa Amicon Ultra-4 cellulose concentrator (cellulose acetate filter). Capsids (50 μM CA) were incubated overnight with 0, 12.5, and 500 μM of LEN at room temperature prior to vitrification in Lacey carbon film 300-mesh copper grids (Electron Microscopy Sciences). Control samples were treated with DMSO.

Projection image data were collected at the University of Utah Electron Microscopy Core facility, as described below. Quantifications of curvature and fracturing were performed as described above.

### Time-of-addition infectivity assays

THP-1 cells (100,000/well) were seeded in 96-well plates, in the presence of 5 μg/mL polybrene and 25 nM (in terms of p24) of VSV-G pseudotyped HIV-GFP virus, in 190 μL of complete media (RPMI, 10% FBS, Pen-Strep). LEN was added to final concentrations of 5 or 250 nM at indicated times post-transduction (**Fig. 5A**). At 24 hours post-transduction, cells were washed, fixed with Cytofix fixation buffer (BD Biosciences), and stained with LIVE/DEAD Fixable Violet stain (405 nm) (ThermoFisher), following the manufacturer’s instructions. Infectivity was assessed by flow cytometry using a FACSCanto (BD Biosciences), acquiring 10,000 events.

### Sample preparation and collection of untilted projection images

LEN-saturated cores used for structure determination were at 5 μM CA and incubated with 50 μM LEN for 1 h and then vitrified in Quantifoil grids, using a Leica GP2 plunge-freezer with the following settings: temperature = 11 °C, humidity = 85%, blot time = 4.5 s. Image data were collected at the University of Utah Electron Microscopy Core facility, using a Krios (ThermoFisher) operating at 300 kV and equipped with an energy filter and K3 direct detector (Gatan). Data (12,050 movies) were collected using EPU (ThermoFisher) with a total dose of approximately 50 electrons/Å^2^ over 40 frames and target defocus of 1.0 to 2.5 μm.

Unbound cores used for structure determination were concentrated to 13 μM CA. Five μL was applied to C-Flat grids and then vitrified. Image data were collected at the University of Virginia Molecular Electron Microscopy Core facility, using a Krios with energy filter and K3 detector. Data (6,065 movies) were collected using EPU with a pixel size of 1.08 Å, total dose of approximately 50 electrons/Å^2^ over 40 frames, and target defocus of 1.0 to 2.5 μm.

Unsaturated cores used for structure determination were at 6.5 μM CA and incubated with 1.625 μM LEN (4:1 CA:LEN ratio) for 1 h at room temperature. The sample was supplemented with 0.01% (w/v) octylglucoside prior to vitrification in C-Flat grids. Image data (12,316 movies) were collected at the University of Virginia as described for unbound cores.

Unsaturated synthetic capsids were at 50 μM CA and incubated with 12.5 μM LEN (4:1 CA:LEN ratio) overnight at room temperature prior to vitrification in Lacey grids. Data (9,183 movies) were collected at University of Utah as above.

### Single-particle averaging map calculations

All image processing, including motion correction, CTF estimations, and map calculations were performed using cryoSPARC (*40*). Particle picking was performed as described (*14*) or by using Topaz (*41*) after training with manual picks. Multiple rounds of reference-free, two-dimensional (2D) classification were performed to identify and discard junk particles, yielding the initial particle sets for each data set.

For the LEN-saturated hexamer, the initial set consisted of 2,155,205 particles. These were further classified through one round of heterogeneous refinement with C6 symmetry imposed, using 8 copies of EMD-3465 as initial templates. The two best classes, containing 997,368 and 304,845 particles, were combined and processed through one round each of homogeneous and local refinement jobs. After discarding duplicates, the final set of 1,064,761 particles was locally refined to yield the final map, which had a nominal resolution of 2.8 Å at the 0.143 FSC cut-off. For focused reconstruction centered on two hexamers, the above map was re-centered and re-aligned with C2 symmetry imposed. Local refinement yielded a map with nominal resolution of 3.2 Å.

For the unbound hexamer, the initial set consisted of 3,339,201 particles. These were further classified through one round of heterogeneous refinement in C1 (no symmetry imposed), using 6 copies of EMD-10226 (the warped hexamer) (*20*) as initial templates. The two best classes (1,259,785 and 707,850 particles) were further cleaned up using one round of 2D classification, yielding 1,199,079 particles that were refined against EMD-10226 in C1. Duplicate removal resulted in a final set of 901,622 particles. The same set was then processed through two parallel reconstruction workflows (in C1 or C6 symmetry) consisting of homogenous and local refinement jobs. The final maps had nominal resolutions of 3.3 Å (C1) and 2.9 Å (C6).

For unsaturated cores, the initial set consisted of 1,599,526 particles. These were further classified through one round of heterogeneous refinement in C1 (no symmetry imposed), using as initial templates 6 copies of a map centered on the pseudo 3-fold connecting two hexamers and one pentamer. The pentamer-containing class (206,965 particles) was subjected to another round of heterogeneous refinement with C5 symmetry imposed and using a re-centered map as template. The final set of 119,349 particles were refined with C5 symmetry, yielding a map with nominal resolution of 4.5 Å. Due to the limited resolution and particle numbers, this data set was not processed further.

For unsaturated in vitro assembled capsids, the initial set consisted of 4,546,022 particles. To reconstruct maps focused on features of the declination, the particles were classified through one round of heterogeneous refinement in C1, using as initial templates 8 copies of a map centered on the pseudo 3-fold connecting two hexamers and one pentamer. After removing duplicates, the remaining 877,757 particles were refined in C1 to yield the final hexamer-hexamer-pentamer map at 3.5 Å resolution. To obtain a map focused on the pentamer, the same particles were refined against a re-centered map, with C5 symmetry imposed. After further duplicate removal, the remaining 518,565 particles were locally refined in C5 symmetry to yield the final pentamer-focused map at 3.5 Å resolution. To obtain a map focused on the hexamer next to the pentamer, particles used for the pentamer reconstruction were subjected to symmetry expansion and refined in C1 against a re-centered map. After another round of duplicate removal, the final set of 1,669,740 particles were locally refined to 3.4 Å nominal resolution. To obtain a map centered on a hexamer-hexamer interface from the main body of the capsid, the initial particle set was classified using a heterogeneous refinement run, using 8 copies of EMDB-3465 as initial templates and with C6 symmetry imposed. Two classes (1,416,153 and 479,318 particles) were selected and combined for further processing. After removing duplicates, the remaining 740,861 particles were first refined in C6, then in C2 after map re-centering. Local refinement yielded a consensus C2 map containing two hexamers at 3.8 Å nominal resolution. The high tilt and low tilt particle subsets were identified and reconstructed with imposed C2 symmetry, following the 3D variability-based tilt and twist analysis approach described below, yielding two maps at 4.1 Å nominal resolution.

Image processing workflows are diagrammed in **Fig. S4** (LEN-saturated cores), **Fig. S5** (unbound cores), **Fig. S8** (unsaturated cores), **Fig. S9** (unsaturated in vitro capsids, declination), **Fig. S11** (unsaturated in vitro capsids, hexamer-hexamer interface), and statistics are in **Table 1**.

### Coordinate modeling

Atomic coordinates were modeled into the unbound (both C1 and C6) and LEN-saturated (C6) maps. Initially, the appropriate number of copies of a hexamer subunit (8ckv) (*15*) were docked into each map. After one or two rigid-body refinement rounds (treating each domain independently), iterative rounds of real-space refinement using the Phenix suite (*42*) and model building using Coot (*43*) were performed. Secondary structure, reference model restraints, and non-crystallographic symmetry (if appropriate) were used. Model quality was continuously monitored using MolProbity (*44*), as implemented in Phenix and Coot. Statistics of the coordinate models are in **Table 2**.

**Table 2.** Coordinate modeling and real space refinement.

|  | Bound Hexamer<br>C6 | Unbound Hexamer<br>C1 | Unbound Hexamer<br>C6 |
| --- | --- | --- | --- |
| EMDB | 72657 | 71816 | 71817 |
| PDB | 9Y7J | 9PRY | 9PRZ |
| Model resolution at FSC 0.143/0.5 threshold (Å) | 2.8/3.0 | 3.2/3.4 | 2.9/3.1 |
| No. of nonhydrogen atoms | 1,726 | 9,691 | 1,615 |
| No. of protein residues | 212 | 1,242 | 206 |
| Protein B-factors (Å<sup>2</sup>) | 67.73 | 68.9 | 53.4 |
| LEN B-factors (Å<sup>2</sup>) | 46.3 | n/a | n/a |
| RMSD bond lengths (Å) | 0.002 | 0.003 | 0.002 |
| RMSD bond angles (°) | 0.437 | 0.458 | 0.404 |
| MolProbity score | 1.11 | 1.14 | 1.03 |
| Clash score | 2.92 | 3.51 | 2.47 |
| Poor rotamers (%) | 1.09 | 0.76 | 0 |
| Ramachandran favored (%) | 99.0 | 98.7 | 99.5 |
| Ramachandran outliers (%) | 0 | 0 | 0 |
| Ramachandran z-score | 2.18 | 1.96 | 1.04 |

### Local curvature analysis of dihexamers

Tilt and twist analysis was performed by adapting a previously described strategy (*15*). 3D variability analysis was performed in cryoSPARC (*45*), starting with maps focused on the hexamer-hexamer interface and a mask covering both hexamers. Components corresponding to tilt and twist were identified by visual inspection. Non-overlapping particle subsets were subsequently defined, seven for each component. The first, fourth, and seventh subsets from each component were pooled in all possible combinations, then subjected to local refinement in C2 symmetry. Statistics of resulting maps are in **Table 3**. We note that this approach underestimates the full range of tilts and twists present in the samples, but sufficient to describe the key differences between the unbound and LEN-bound capsids. Tilt and twist angles were measured as described (*15*), except that PyMol (Schrodinger Scientific) instead of Chimera software was used to measure angles.

**Table 3.**
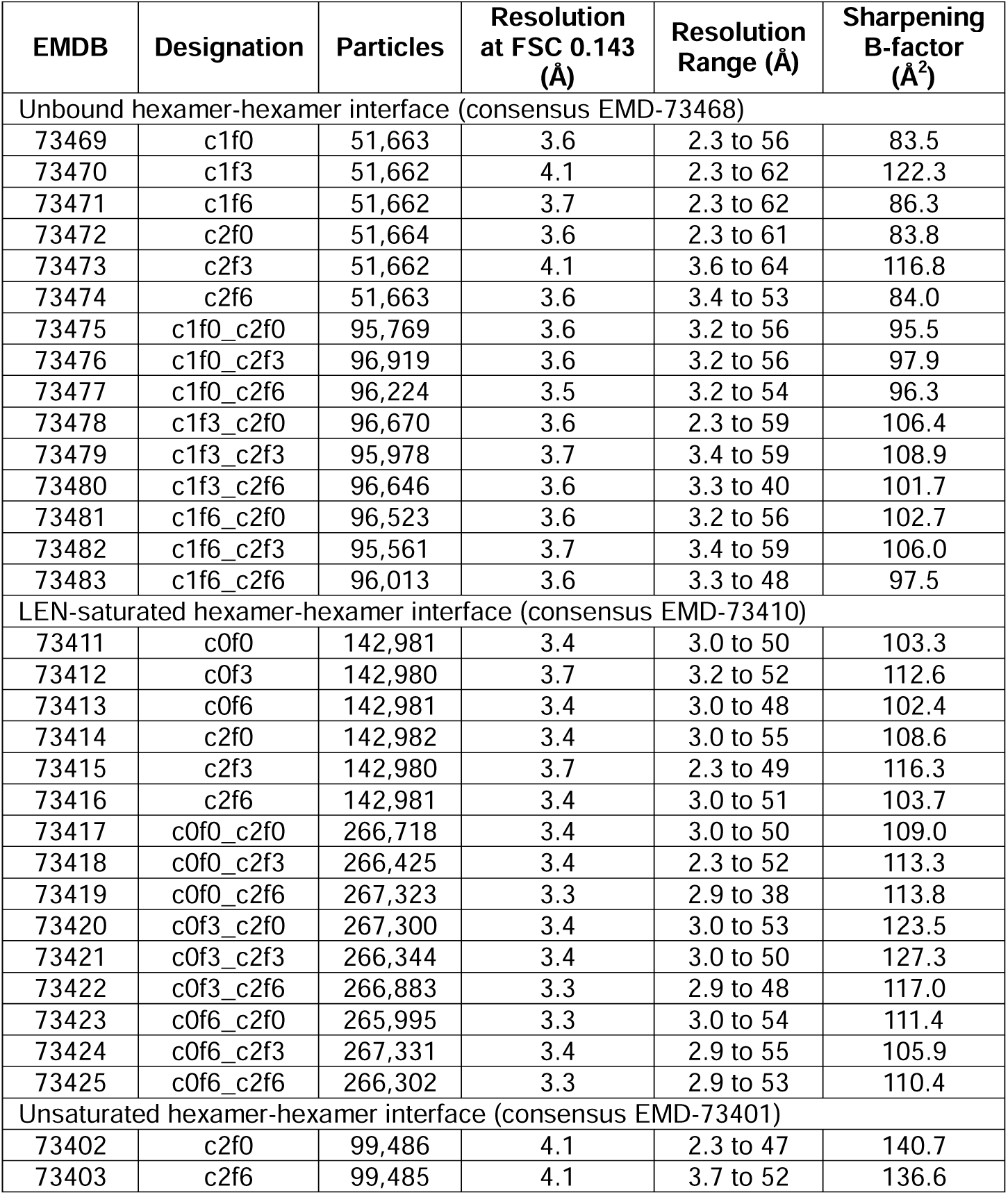
Reconstructions of hexamer-hexamer interface tilt/twist subpopulations.

### Cryotomography analysis

LEN-saturated cores were prepared as above for untilted data collection, and then mixed with equal volume of 10-nm BSA Gold Tracer (Electron Microscopy Sciences), prior to vitrification in C-flat 2/1 300-mesh grids. Cryotomograms were acquired using the University of Utah Electron Microscopy Core facility Krios microscope. Tilt series were collected using the data collection software SerialEM (*46*) with an angular range of −60° to +60°, angular increment of 2°, target defocus value of 4 μm, and a nominal magnification of ×33,000, which corresponds to a pixel size of 2.68 Å. Images were aligned by using IMOD (*47*). Lattice mapping – using the Dynamo software package (*48*) – and lattice map visualization were performed as described (*32, 49*).

### Depositions

The following maps have been deposited at the EMDB: unbound hexamer C1 symmetry, EMD-71816; unbound hexamer C6 symmetry, EMD-71817; unbound hexamer-hexamer interface consensus map, EMD-73468, and tilt/twist series from particle subsets of this map, EMD-73469 through EMD-73483; saturated hexamer C6 symmetry, EMD-72657; saturated hexamer-hexamer interface consensus map, EMD-73410 and tilt/twist series from particle subsets of this map, EMD-73411 through EMD-73425; pentamer from unsaturated cores, EMD-73397; hexamer-hexamer-pentamer pseudo 3-fold focused map from unsaturated capsids, EMD-73398; pentamer from unsaturated capsids, EMD-73399; hexamer next to pentamer from unsaturated capsids, EMD-73400; hexamer-hexamer interface from unsaturated capsids consensus map, EMD-73401, low-tilt subset, EMD-73402, and high-tilt subset, EMD-73403. Coordinate models were deposited at the PDB: unbound hexamer C1 symmetry, 9pry; unbound hexamer C6 symmetry, 9prz; saturated hexamer C6 symmetry, 9y7j.

## Acknowledgements

We thank T. Porter for assistance with protein purification, M. Purdy for expert support with image data collection, and J. Perilla for useful discussions. We thank Gilead Sciences for providing LEN at early stages of the project. This study was funded by NIH grant U54-AI170856. O.P. and N.F.B.d.S. were also supported by R37-AI150479. J.A.L. was supported by F32-AI195016.

