## Supplementary Figures for "Lenacapavir allosterically remodels the HIV-1 capsid"

Figure S1 – Purification and LEN treatment of HIV-1 cores
Figure S2 – LEN treatment of cores and synthetic capsids at high concentrations
Figure S3 – Cryotomography of LEN-fractured cores
Figure S4 – Focused reconstructions of LEN-saturated CA hexamer
Figure S5 – Focused reconstructions of unbound CA hexamer
Figure S6 – Structural comparisons
Figure S7 – Gallery of maps from particle subsets of unbound and bound hexamer-hexamer interfaces
Figure S8 – Focused reconstruction from cores incubated with sub-saturating amounts of LEN (4:1 CA to drug ratio)
Figure S9 – Focused reconstructions on declinations from in vitro assembled, empty capsids incubated with sub-saturating amounts of LEN (4:1 CA to drug ratio)
Figure S10 – LEN occupancies in the independent hexamers from the map centered on the hexamer-hexamer-pentamer pseudo 3-fold
Figure S11 – Focused reconstructions on the hexamer-hexamer interface from in vitro assembled, empty capsids incubated with sub-saturating amounts of LEN (4:1 CA to drug ratio)

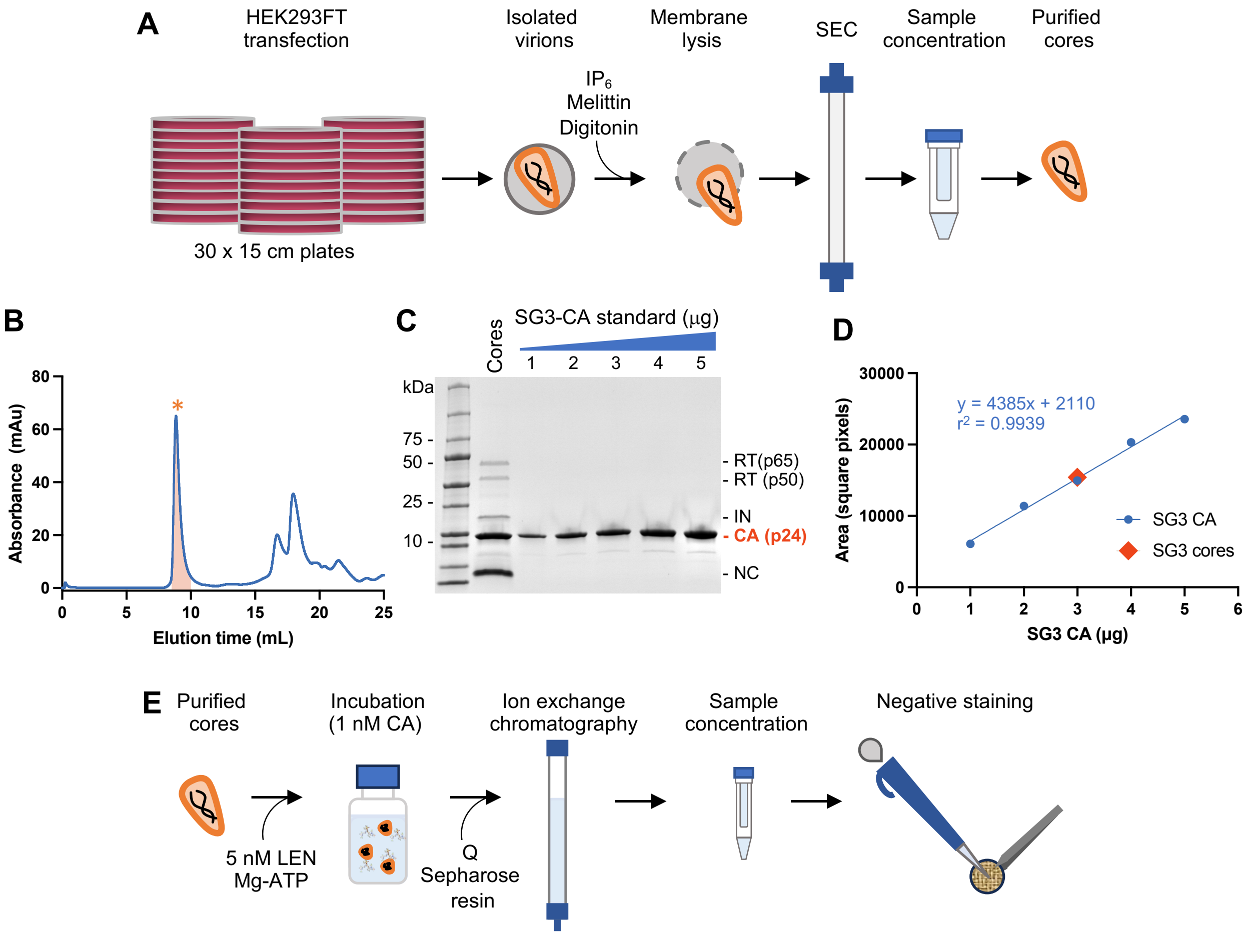


**Figure S1.** Purification and LEN treatment of HIV-1 cores. (**A**) Protocol outline for core purification. (**B**) Elution profile of lysed virions on a Superdex 200 size exclusion column. Asterisk indicates core peak. (**C**) Coomassie-stained SDS-PAGE gel of purified cores. Lanes 1-5 contain known amounts of purified recombinant CA protein as quantification standards. RT – reverse transcriptase, IN – integrase, CA – capsid protein, NC – nucleocapsid protein. (**D**) Densitometry quantification of the gel in C. Blue circles correspond to points on the standard curve. Orange diamond indicates the concentration of purified cores (expressed as CA concentration). (**E**) Schematic protocol of treatment with 5 nM LEN. Cores (equivalent to 1 nM of CA) were incubated with LEN or DMSO control.

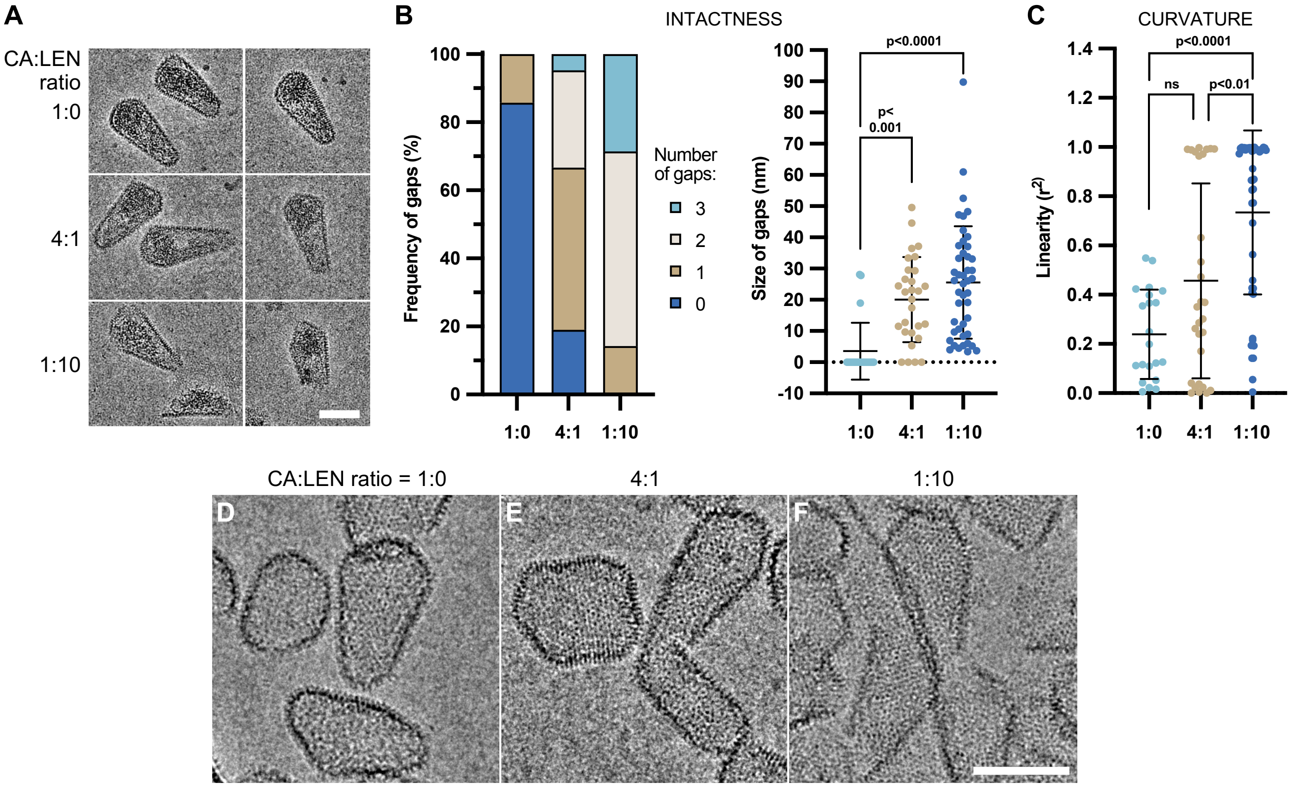


**Figure S2.** LEN treatment of cores and synthetic capsids at high concentrations. (**A**) CryoEM images of representative cores (~5 μM CA) after incubation with indicated amount of LEN. Scale bar = 50 nm. (**B**) Quantification of fracturing (n = 21 randomly selected cores for each condition). Left panel, number of gaps observed per capsid. Right panel, sizes of the gaps. (**C**) Quantification of curvature for the same cores. (**D-F**) CryoEM images of in vitro assembled, empty capsids incubated with the indicated amounts of LEN. Scale bar = 50 nm.

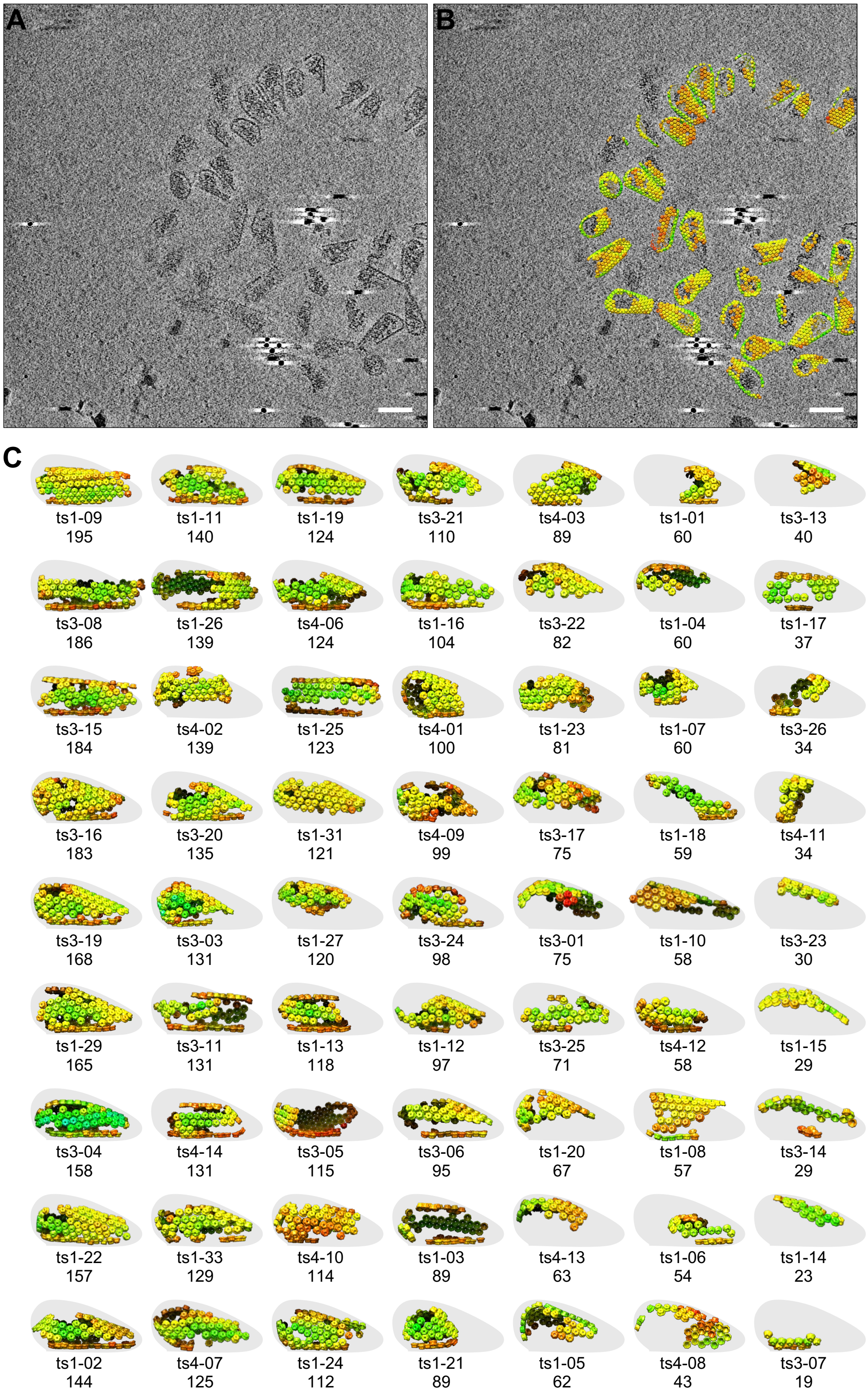


**Figure S3.** Cryotomography of LEN-fractured cores. (**A,B**) Slice of representative cryotomogram, without (A) and with (B) lattice maps overlaid. Scale bars, 100 nm. (**C**) Gallery of lattice maps from 63 cores that could be identified as having capsid remnants with similar characteristics. Each lattice map is oriented with the flat walls (or deduced flat wall) at the bottom. The general cone shape is outlined in gray.

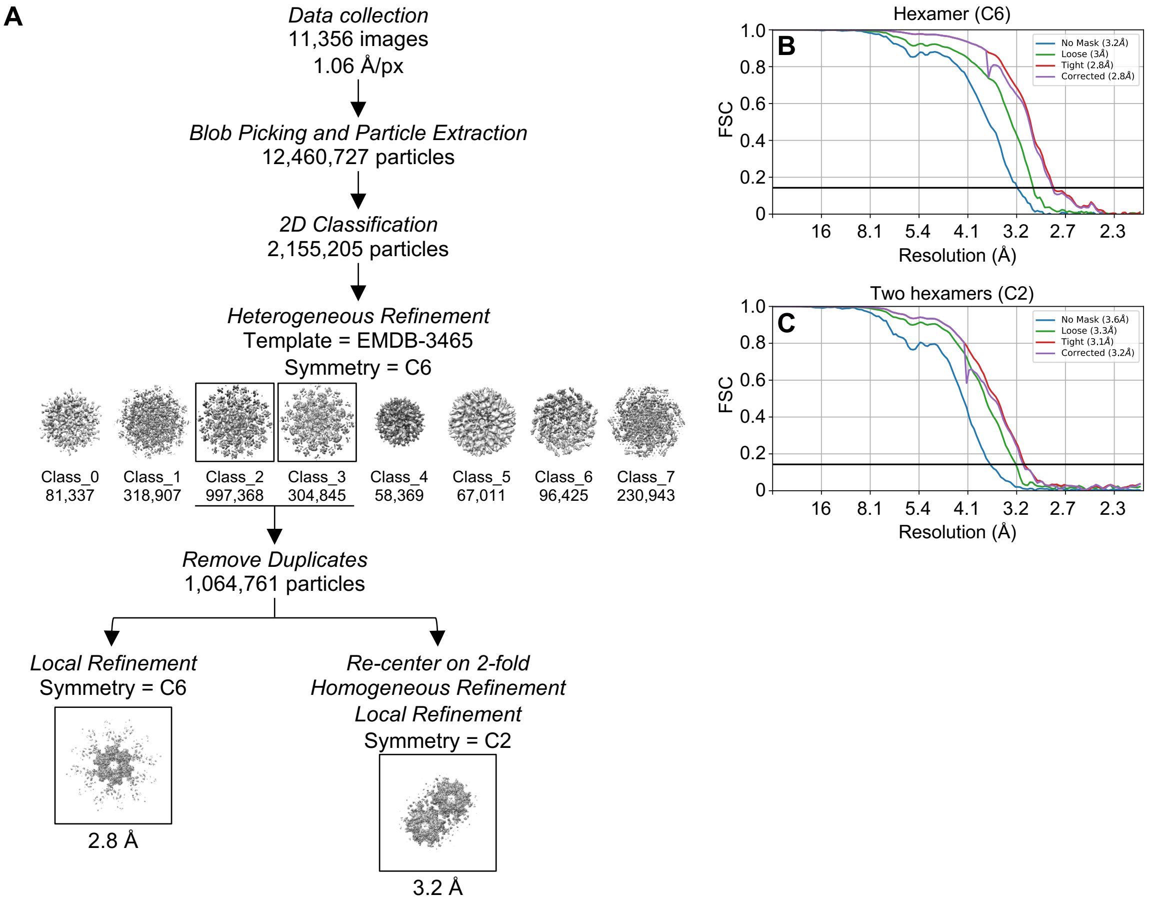


**Figure S4.** Focused reconstructions of LEN-saturated CA hexamer. (**A**) CryoEM workflow. (**B**) Fourier shell correlation curves for the indicated maps.

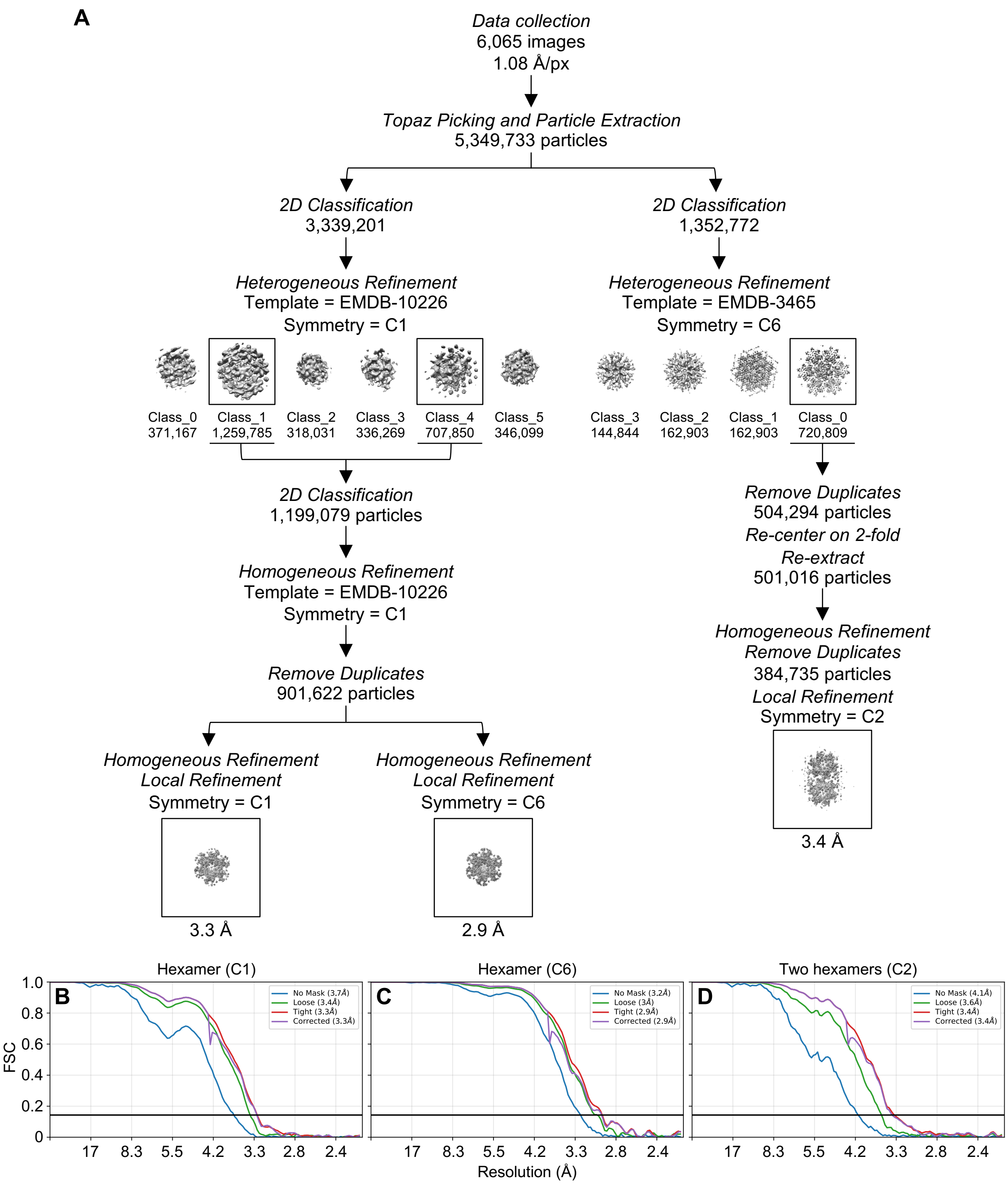


**Figure S5.** Focused reconstructions of unbound CA hexamer. (**A**) CryoEM workflow. (**B**) Fourier shell correlation curves for the indicated maps.

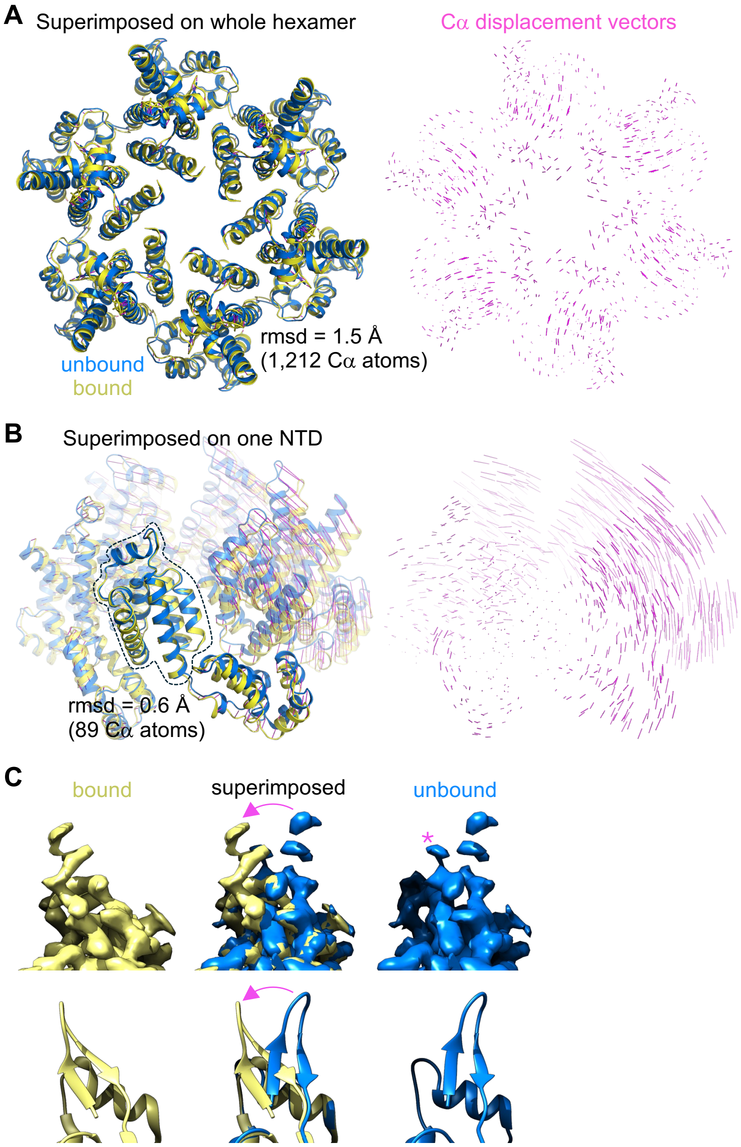


**Figure S6.** Structural comparisons. (**A**) Superposition of unbound (blue) and LEN-bound (yellow) structures as whole hexamer units, with displacement vectors on the right (magenta). (**B**) Superposition on indicated NTD alone. (**C**) Altered configuration of the N-terminal β-hairpin. Arrows indicate movement from unbound to bound configurations. Asterisk indicates unmodeled density, suggesting that the bound configuration (or similar) may represent a minor population in the unbound map.

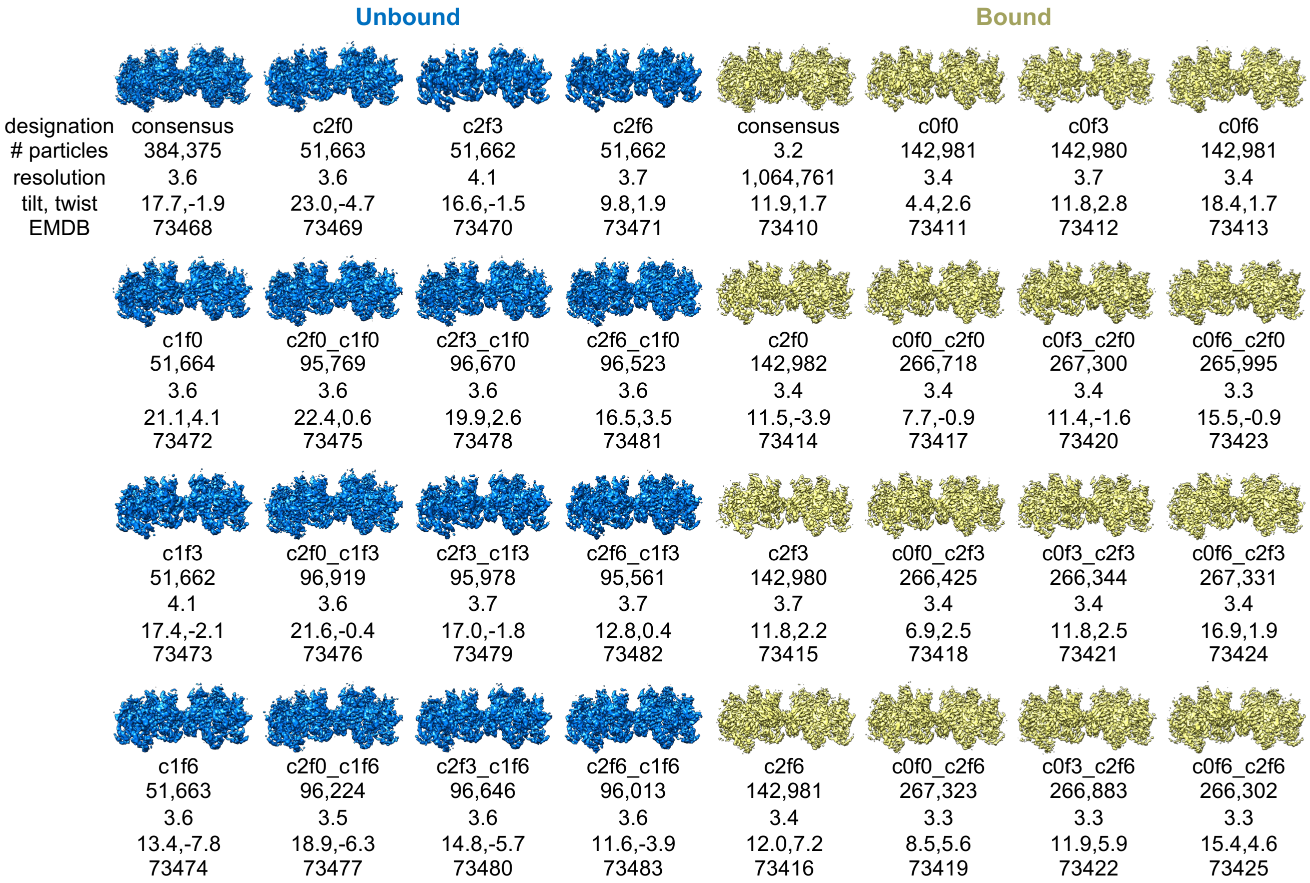


**Figure S7.** Gallery of maps from particle subsets of unbound (left) and bound (right) hexamer-hexamer interfaces. Map designations, number of particles, nominal resolutions after local refinement, as well as measured tilt and twist angles, are indicated.

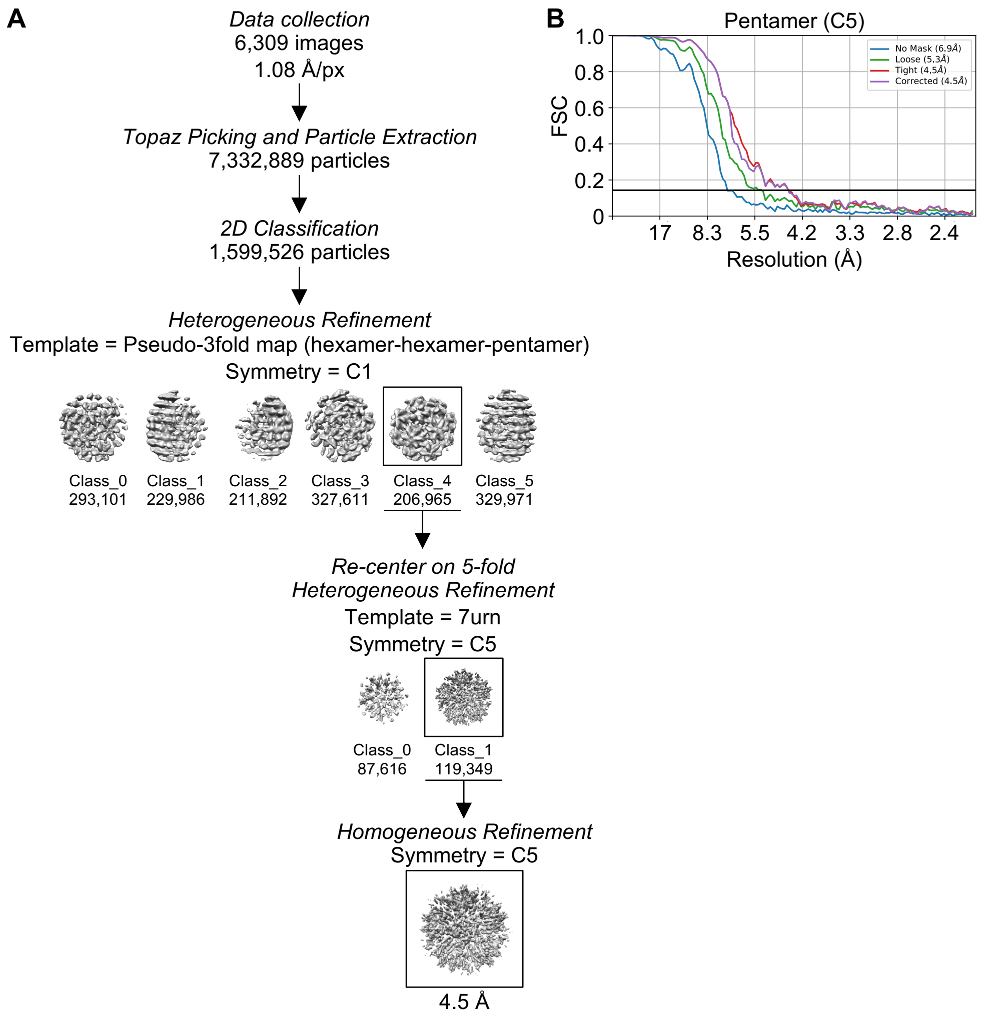


**Figure S8.** Focused reconstruction from cores incubated with sub-saturating amounts of LEN (4:1 CA to drug ratio). (**A**) CryoEM workflow. (**B**) Fourier shell correlation curves for the final pentamer-focused map. Due to limited resolution and particle numbers, further processing was not pursued for this data set.

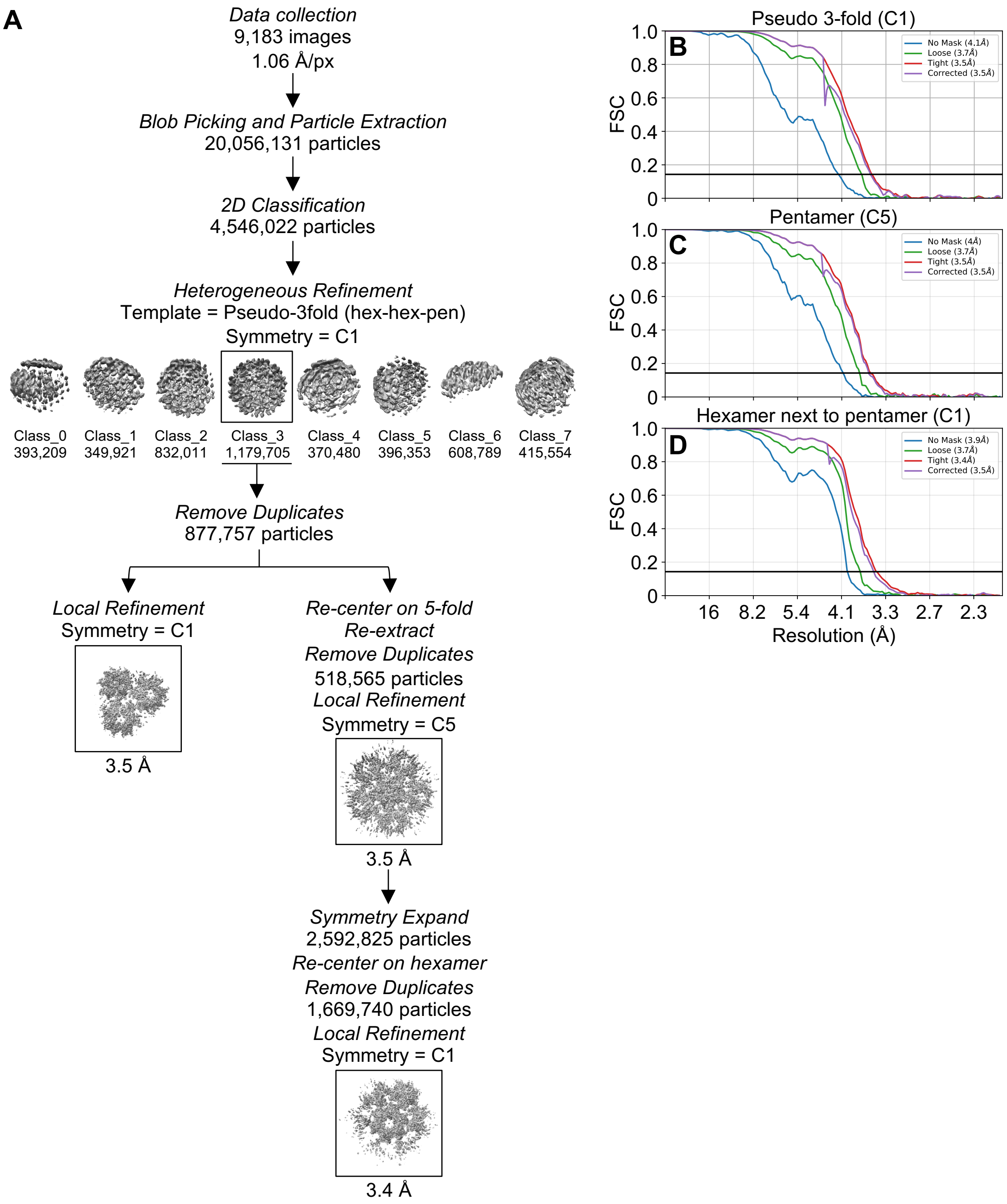


**Figure S9.** Focused reconstructions on declinations from in vitro assembled, empty capsids incubated with sub-saturating amounts of LEN (4:1 CA to drug ratio). (**A**) CryoEM workflow. (**B-D**) Fourier shell correlation curves for the indicated maps.

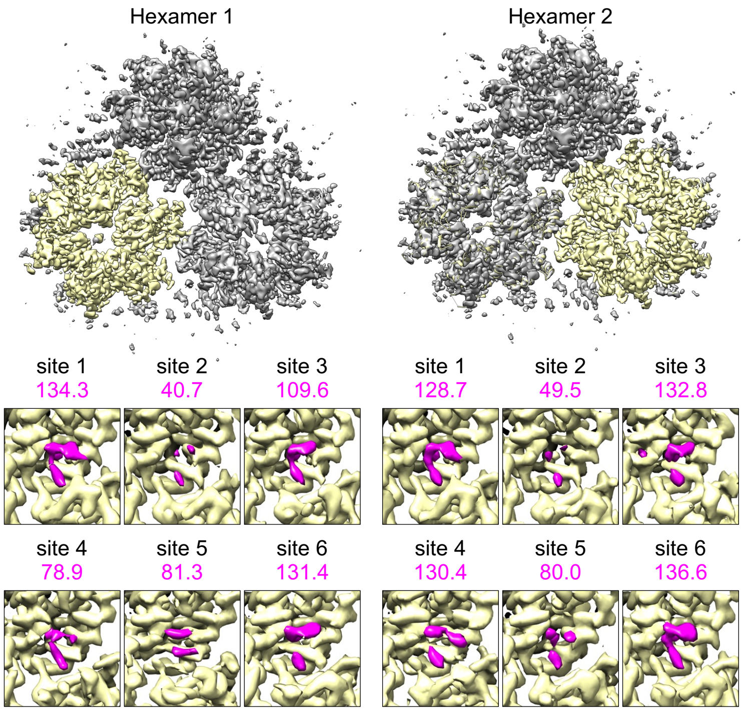


**Figure S10.** LEN occupancies in two independent hexamers from a map centered on the hexamer-hexamer-pentamer pseudo 3-fold. Numbers in magenta indicate the measured volumes ($Å^3$) of drug densities at map contour level of 0.24.

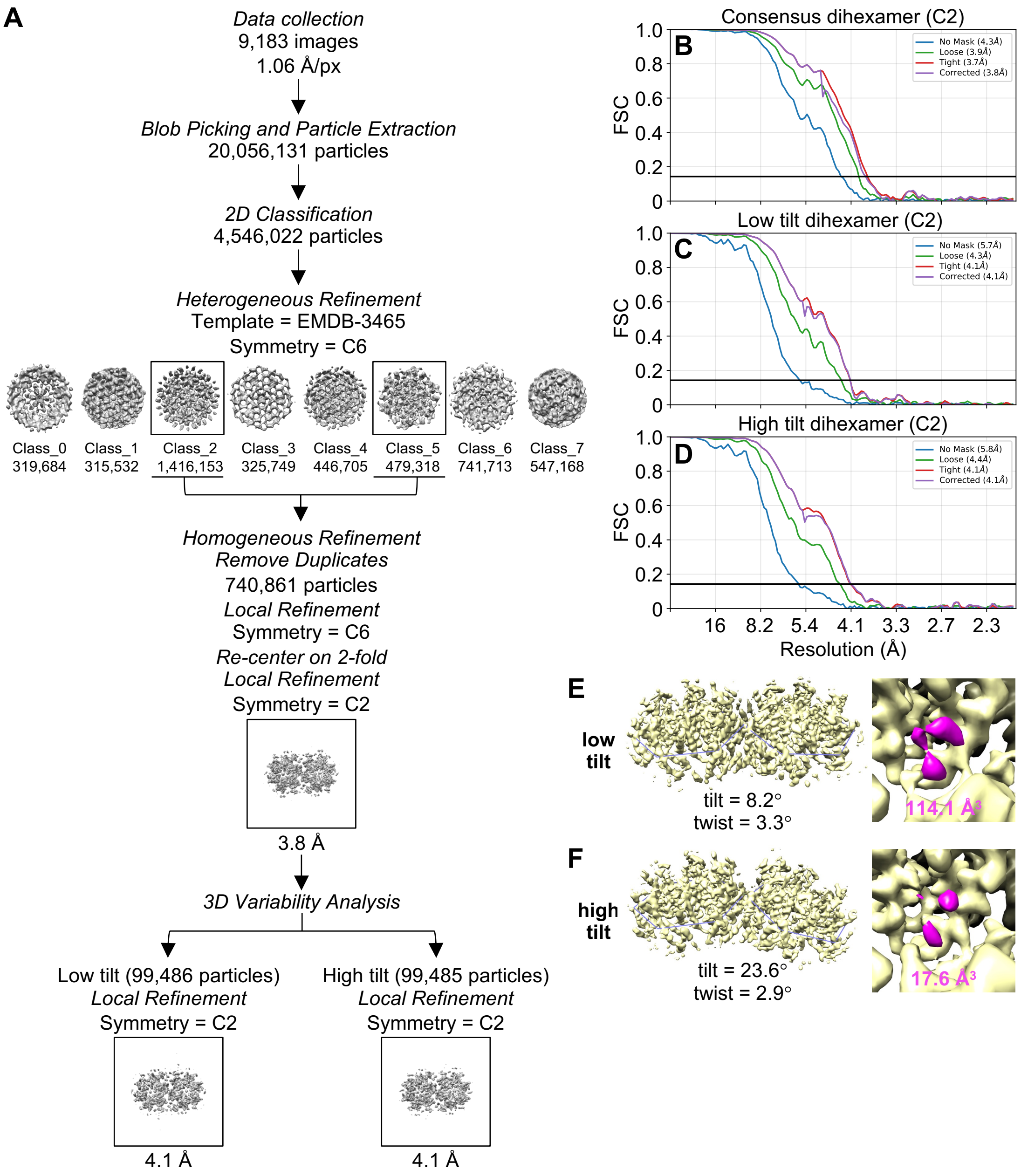


**Figure S11**. Focused reconstructions on the hexamer-hexamer interface from in vitro assembled, empty capsids incubated with sub-saturating amounts of LEN (4:1 CA to drug ratio). (**A**) CryoEM workflow. (**B-D**) Fourier shell correlation curves for the indicated maps. (**E-F**) LEN occupancies expressed as blob volume ($Å^3$) measured at map contour level of 0.19, from low tilt (E) and high tilt (F) hexamer-hexamer interfaces.
